# Iron Deficiency Drives Sarcopenia in the Elderly: HIF-1α-Mediated Fibro-Adipogenic Progenitor Differentiation Induces Fat Infiltration and Impairs Muscle Function

**DOI:** 10.64898/2026.01.15.699636

**Authors:** Quanzhong Ren, Guihe Yang, Dingding Wang, Wenkai Wu, Yi Wang, Jiahua Feng, Kairui Ma, Anyi Guo, Mingxing Fan, Yuqing Sun, Zhao Lang, Xieyuan Jiang, Yajun Liu, Ling Wang, Renxian Wang

## Abstract

Abnormal iron metabolism is closely linked to sarcopenia; however, the specific iron metabolism features of fat-infiltrating sarcopenia remain poorly understood. Proteomic sequencing revealed that in skeletal muscle with severe fat infiltration, the expression of iron utilization-related proteins was significantly downregulated, whereas that of iron uptake and storage proteins was markedly upregulated, thereby presenting an abnormal iron deficiency (ID) phenotype. Nevertheless, the mechanism by which ID drives intramuscular fat infiltration has not been fully elucidated. Using clinical samples, aged murine ID models, and in vitro cell assays, this study is the first to demonstrate that ID stabilizes hypoxia-inducible factor-1α (HIF-1α), promoting aberrant adipogenic differentiation of fibro-adipogenic progenitors (FAPs), disrupting the homeostatic balance between satellite cells (SCs) and FAPs, exacerbating skeletal muscle fat infiltration, and impairing muscle repair capacity. Notably, treatment with the HIF-1α inhibitor PX478 reversed these pathological alterations and improved muscle function. Collectively, our findings identify the ID-HIF-1α-FAPs axis as a key driver of intramuscular fat infiltration, offering a novel therapeutic target for the clinical management of fat-infiltrating sarcopenia.

## 1. Introduction

Iron is an indispensable micronutrient for various physiological processes of skeletal muscle, and its core functions run through key links such as oxygen transport, mitochondrial respiration, and cellular metabolism.^[1–4]^ Imbalance of iron homeostasis is closely associated with a variety of muscle diseases, among which iron overload and ferroptosis have garnered widespread attention;^[5–7]^ meanwhile, iron deficiency (ID), a condition characterized by insufficient dietary intake or low systemic iron utilization efficiency that fails to meet physiological needs, cannot be ignored.^[8,9]^ ID has a high incidence in the elderly population, with inducing factors including inadequate dietary iron intake, intestinal absorption disorders, and increased metabolic demands of the body.^[10,11]^

In skeletal muscle, ID can lead to impaired muscle function, decreased exercise capacity, and disrupted cellular metabolism. These pathological defects are believed to originate from the adaptive response of cells to iron deprivation, which may involve abnormal changes in mitochondrial biogenesis, energy metabolism, and stress response pathways.^[11–14]^ Notably, recent studies have shown that ID may also disrupt the local cellular microenvironment, thereby affecting the function of satellite cells (SCs) that are crucial for muscle repair and homeostasis maintenance.^[15]^ Skeletal muscle repair is a sophisticated, dynamic process orchestrated by the synergistic interplay of multiple cell populations, including SCs, fibro-adipogenic progenitors (FAPs), and immune cells.^[15,16]^ Among these cellular components, the functional equilibrium between SCs and FAPs constitutes the cornerstone of skeletal muscle homeostasis: SCs initiate and drive the entire cascade of muscle regeneration, whereas FAPs support tissue repair via paracrine actions but tend to undergo aberrant adipogenic differentiation under pathological conditions,^[17]^ ultimately leading to intramyocellular fat infiltration (IMFI), which is a canonical pathological hallmark of sarcopenia.^[18]^ Notably, a study has confirmed that inhibition of ferroptosis facilitates the adipogenic differentiation of FAPs.^[19]^ This finding suggests that FAPs are a cell type highly sensitive to iron status. Yet, the impacts of iron metabolic disorders, particularly ID, on FAPs remain largely elusive.

Furthermore, hypoxia-inducible factor 1α (HIF-1α), as a core molecule regulating cellular adaptation to hypoxia and metabolic stress,^[20–24]^ its functional regulation is closely related to iron metabolism and adipogenesis. Under iron-deficient conditions, the activity of prolyl hydroxylase domain 2 (PHD2), which is responsible for targeted degradation of HIF-1α, is inhibited, resulting in a significant enhancement of HIF-1α protein stability.^[25]^ Existing studies have confirmed that high levels of HIF-1α can promote adipogenic differentiation of various cells,^[26–29]^ suggesting that there may be an inherent regulatory association between ID, HIF-1α stabilization, and FAPs-mediated intramyocellular fat accumulation. However, although the above association has been initially verified, the specific molecular mechanisms by which ID induces muscle dysfunction, especially the synergistic regulatory role of FAPs and the HIF-1α signaling pathway, remain incompletely elucidated.

In this study, we hypothesized that ID-induced HIF-1α activation drives adipogenic differentiation of FAPs, thereby promoting fatty infiltration and impairing muscle repair. To test this hypothesis, we integrated analyses of clinical human samples, a mouse model of systemic ID using 18-month-old aged mice and in vitro FAP cell experiments. Specifically, we aimed to: (1) characterize the imbalance in iron homeostasis in elderly individuals with high fatty infiltration; (2) validate the causal role of ID in muscle repair dysfunction and fatty infiltration using the aforementioned animal model; (3) explore the transcriptomic and functional alterations in FAPs under ID conditions; (4) identify HIF-1α as a core mediator of FAP dysregulation; and (5) evaluate the therapeutic potential of HIF-1α inhibition in reversing ID-related muscle abnormalities. Our findings provide novel insights into the pathogenesis of age-related sarcopenia complicated by fatty infiltration and offer a promising therapeutic target for clinical intervention.

## 2. Results

### 2.1. Skeletal Muscle with High Fat Infiltration Exhibits Actual ID

To elucidate the mechanistic underpinnings of IMFI in the pathogenesis of sarcopenia, we employed magnetic resonance imaging (MRI) to accurately quantify the intramyocellular fat fraction (IMFF) in the paraspinal muscles of participants. This was coupled with isokinetic strength measurements of trunk muscles to delineate the threshold effect of IMFI on muscle strength (**Fig. 1a**). Based on IMFF levels, participants were stratified into four groups: <5%, 5-10%, 10-15%, and >15% (**Supplementary Fig.1a**). This stratification was guided by quartiles of the population distribution, accounting for sex differences and clinical applicability.

**Fig. 1.**
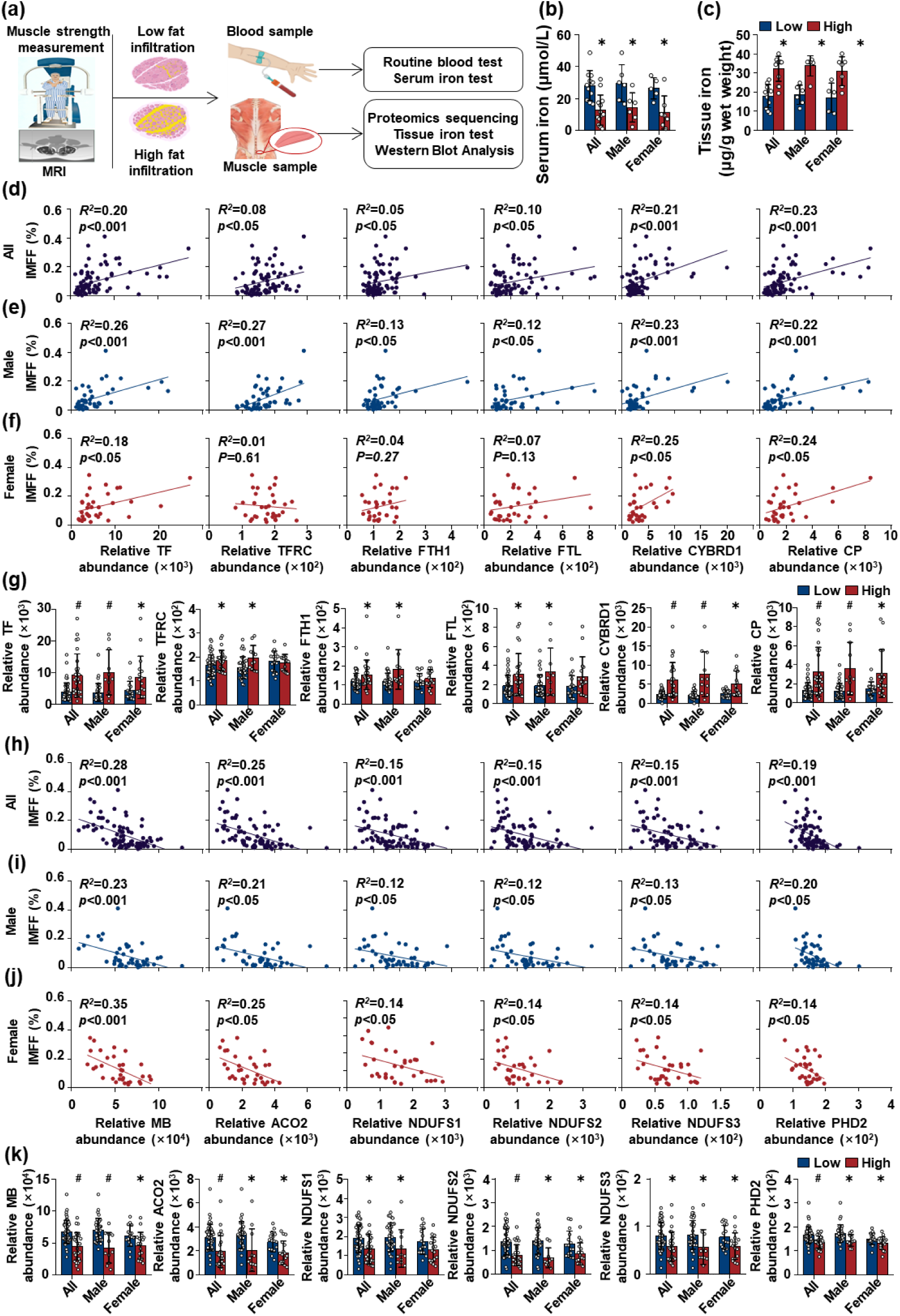
High IMFI is associated with iron metabolism dysregulation. (a) Schematic diagram of the study design: Participants were stratified into Low (IMFF < 5%) and High (IMFF > 15%) groups based on IMFF quantified by MRI. Clinical data, serum iron, tissue iron, isokinetic muscle strength, and proteomic profiles were analyzed to explore the association between iron metabolism and IMFI. (b) Comparison of serum iron levels between Low and High groups in the total population (Low: n=11; High: n=13), males (Low: n=6; High: n=6), and females (Low: n=5; High: n=7). (c) Comparison of Non-heme iron levels between Low and High groups in the total population (Low: n=11; High: n=13), males (Low: n=6; High: n=6), and females (Low: n=5; High: n=7). (d-f) Correlation analyses between iron metabolism-related proteins (TF, TFRC, FTH1, FTL, CYBRD1, CP) and IMFF (%) in the total population (d, Low: n=51; High: n=28), males (e, Low: n=35; High: n=11), and females (f, Low: n=16; High: n=17). Protein expression levels are presented as normalized relative abundance. (g) Protein expression levels of iron metabolism-related proteins in Low vs. High groups, stratified by gender (Low: n=51; High: n=28; males: Low=35, High=11; females: Low=16, High=17). Protein expression levels are presented as normalized relative abundance. (h-j) Correlation analysis between iron utilization-related proteins (MB, ACO2, NDUFS1/2/3, PHD2) and IMFF (%) in the total population (h, Low: n=51; High: n=28), males (e, Low: n=35; High: n=11), and females (f, Low: n=16; High: n=17). Protein expression levels are presented as normalized relative abundance. (k) Protein expression levels of iron utilization-related proteins in Low vs. High groups, stratified by gender (Low: n=51; High: n=28; males: Low=35, High=11; females: Low=16, High=17). Protein expression levels are presented as normalized relative abundance. * indicates a significant difference (p < 0.05) in the High group compared to the Low group; # indicates a significant difference (p < 0.001) in the High group compared to the Low group.

Muscle strength assessments revealed a marked reduction in the paraspinal muscle index (PSMI), a validated measure of muscle mass, when IMFF exceeded 15%. Concomitantly, significant reductions in extension, rotational, and lateral flexion strength were observed in this subgroup (**Supplementary Fig.1b**). Based on these findings, participants with IMFF >15% were defined as the High-IMFF group and those with IMFF <5% as the Low-IMFF group. Hematoxylin and eosin (H&E) staining revealed a marked increase in fat infiltration in the muscle tissues of the High-IMFF group (**Supplementary Fig.2**). The High IMFF group was therefore prioritized for subsequent investigations. Given the established link between iron metabolism disorders and pathological muscle remodeling,^[30–32]^ we next focused on iron metabolism abnormalities to investigate whether they constitute a key pathogenic driver of IMFI development.

**Fig. 2.**
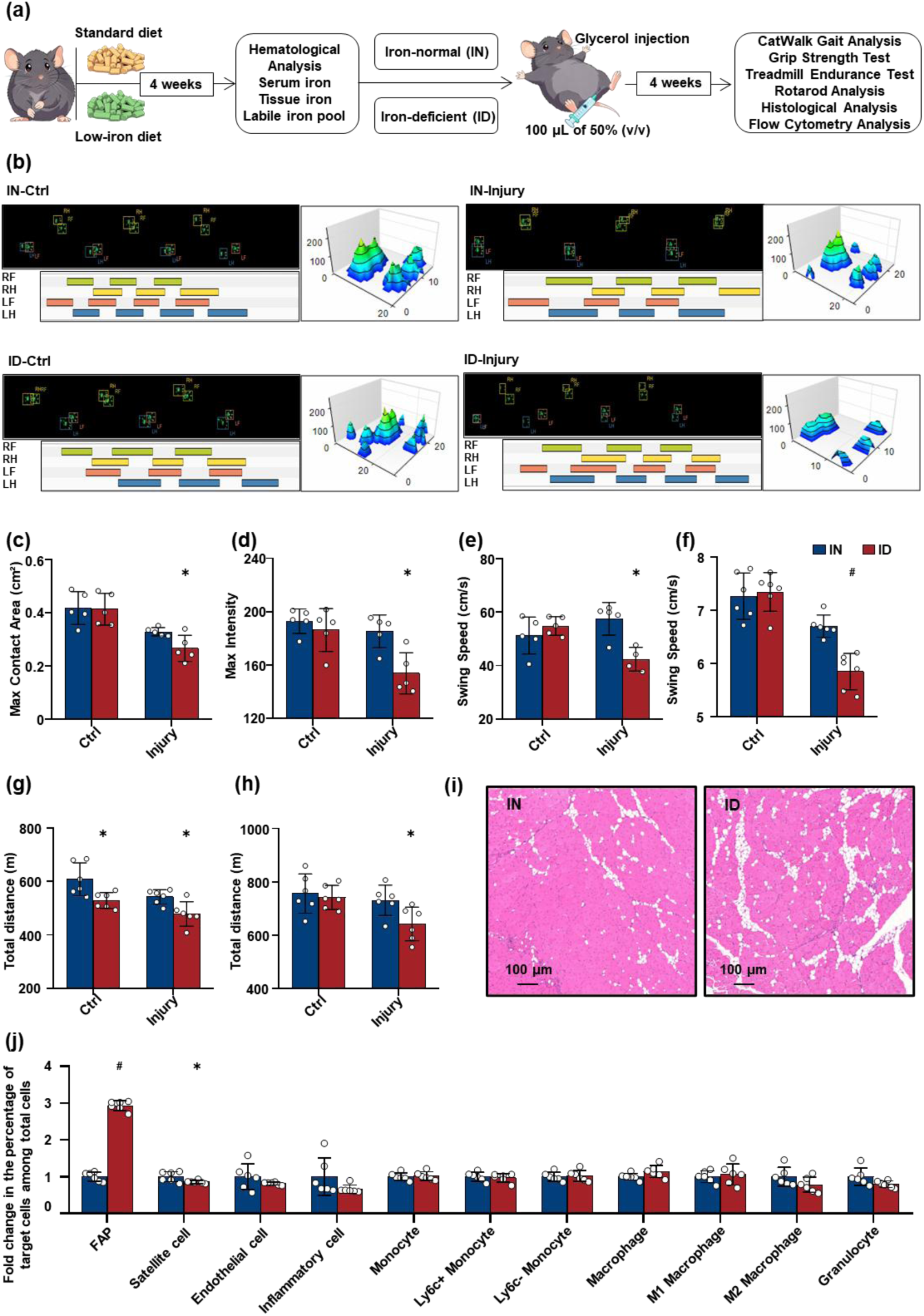
Behavioral, histological, and flow cytometric analyses of muscle function and cellular composition after glycerol-induced injury in IN and ID mouse models. (a) Experimental workflow: Mice were fed either a standard diet or a low-iron diet for 4 weeks. After evaluating iron metabolic indices, muscle injury was induced by intramuscular injection of 100 μL 50% (v/v) glycerol, while control mice received saline injection. Four weeks after modeling, analyses including CatWalk gait analysis, grip strength test, treadmill endurance test, rotarod test, histological examination, and flow cytometry were performed. (b) Visualization of CatWalk gait analysis: Representative hindlimb footprint distributions (top panels), strip plots of gait parameters (bottom panels), and 3D pressure distribution maps of the left hindlimb (right panels) for IN and ID mice in control (Ctrl) and injury groups; distinct colors denote different hindlimbs. (c–e) Gait analysis results: (c) Maximum contact area (cm²), (d) Maximum intensity, and (e) Swing speed (cm/s) in Ctrl and injury groups of IN and ID mouse models (n=5). (f) Maximal grip strength (g/g body weight) in Ctrl (saline-injected) and injury (glycerol-injected) groups of IN and ID mouse models (n=6). (g) Total treadmill exercise distance (m) in Ctrl and injury groups of IN and ID mouse models (n=6). (h) Rotarod retention time (s) in Ctrl and injury groups of IN and ID mouse models (n=6). (i) H&E staining of muscle tissues from Ctrl and Injury groups in IN and ID models. Scale bar = 100 μm. (j) Fold changes in the percentage composition of distinct cell populations in IN and ID groups (n=6). * indicates a significant difference (p < 0.05) in the ID group compared to the IN group; # indicates a significant difference (p < 0.001) in the ID group compared to the IN group.

To elucidate the association between dysregulated skeletal muscle iron levels and fat infiltration, we conducted a comparative analysis of iron-related parameters between the Low and High groups. Hematological analysis revealed no significant differences in anemia-related parameters in the High group, a trend that was consistent in both male and female subgroups (**Supplementary Fig.3**). However, serum iron levels were significantly downregulated, and this trend was consistent across both male and female subgroups (**Fig. 1b**). In contrast, assessment of tissue iron levels demonstrated a significant upregulation of non-heme iron levels in the skeletal muscle of the High group (**Fig. 1c**). This dissociated pattern, characterized by decreased serum iron alongside increased tissue iron, indicates systemic ID and impaired iron metabolic homeostasis within the skeletal muscle. Therefore, dysregulation of iron metabolic homeostasis may be a critical factor contributing to fat infiltration.

**Fig. 3.**
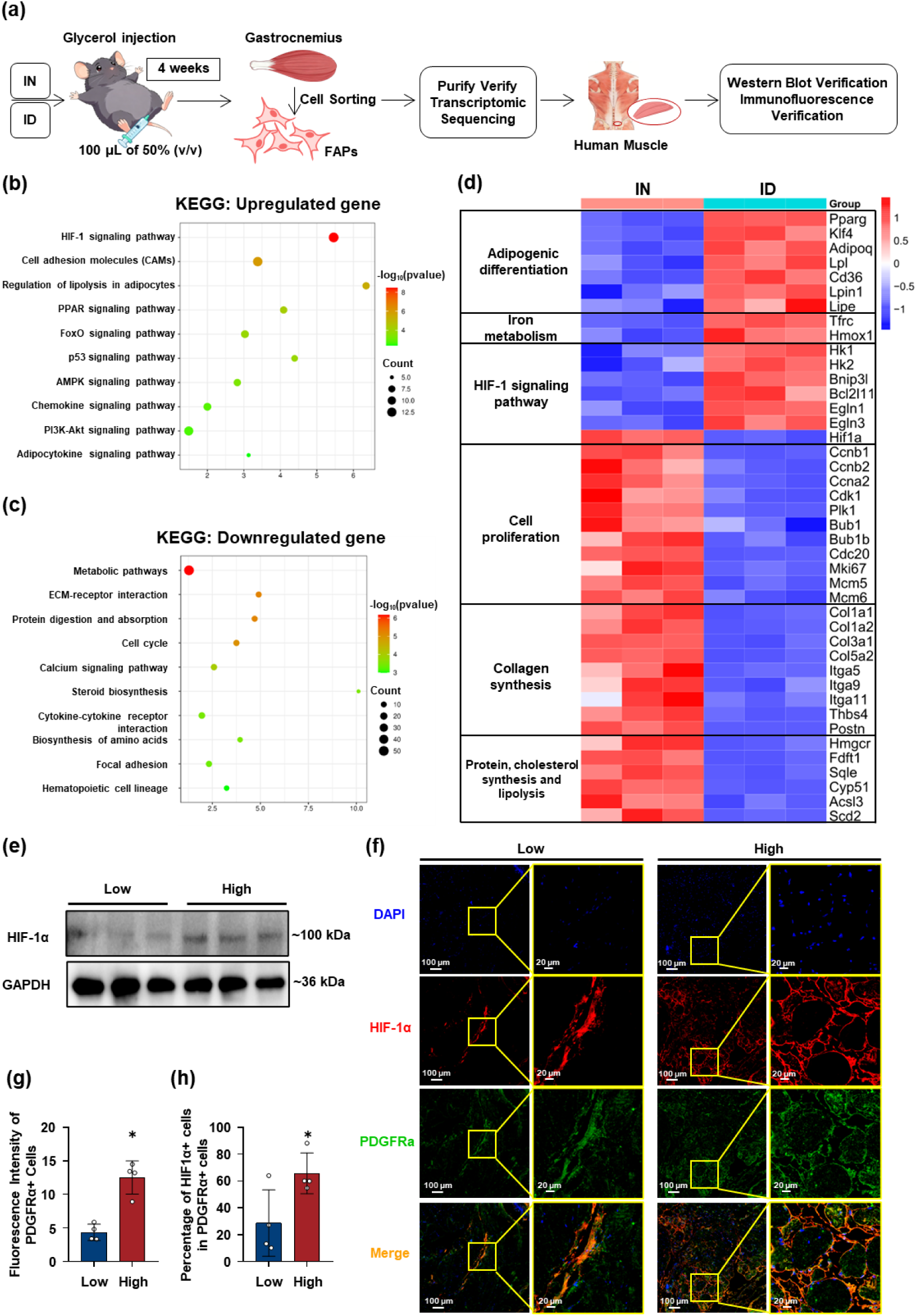
Transcriptomic profiling of FAPs from injured gastrocnemius muscles of IN and ID mice, and characterization of HIF-1α signaling in FAPs (PDGFRα⁺ cells) from clinical skeletal muscle samples with variable fat infiltration. (a) Experimental workflow: IN and ID mice were maintained for 4 weeks, followed by intramuscular injection of 100 μL 50% (v/v) glycerol to induce muscle injury. At 4 weeks post-injury, gastrocnemius muscles were harvested; FAPs were isolated via flow cytometry, with subsequent purity verification and transcriptomic sequencing. Corresponding validation was performed using human skeletal muscle samples. (b) KEGG pathway enrichment bubble plot for upregulated DEGs in ID FAPs (vs. IN FAPs). (c) KEGG pathway enrichment bubble plot for downregulated DEGs in ID FAPs (vs. IN FAPs). (d) Heatmap of gene expression across key functional pathways: Heatmap showing the expression levels of genes involved in adipogenic differentiation, iron metabolism, HIF-1 signaling, cell proliferation, collagen synthesis, and protein/cholesterol synthesis and lipolysis in IN and ID FAPs. (e) Western blot analysis of HIF-1α protein levels in Low and High groups, with Glyceraldehyde-3-phosphate dehydrogenase (GAPDH) as a loading control. (f) Immunofluorescence staining of skeletal muscle sections from Low and High groups, labeled with DAPI (blue, nuclei), HIF-1α (red), PDGFRα (green, marking FAPs), and merged images, with magnified insets. (g) Fluorescence intensity of PDGFRα⁺ FAPs in Low and High groups (n=4). (h) Percentage of HIF-1α⁺ cells among PDGFRα⁺ FAPs in Low and High groups (n=4). * indicates a significant difference (p < 0.05) in the High group compared to the Low group.

To dissect the molecular mechanisms underlying iron metabolic dysfunction, we employed proteomic techniques to examine changes in the expression of iron metabolism-related proteins. Principal Component Analysis (PCA) revealed distinct global protein expression profiles between the Low and High groups (**Supplementary Fig.4a**). Differential protein analysis identified 1809 significantly upregulated proteins and 319 significantly downregulated proteins (**Supplementary Fig.4b**). Kyoto Encyclopedia of Genes and Genomes (KEGG) pathway enrichment analysis indicated that the downregulated proteins were primarily enriched in energy metabolism-related pathways, such as oxidative phosphorylation (**Supplementary Fig.4c**). Conversely, the upregulated proteins were enriched in inflammation and tissue remodeling-related pathways, including complement and coagulation cascades, and ECM-receptor interaction (**Supplementary Fig.4d**). These pathway alterations collectively reveal the pathological characteristics of skeletal muscle with high fat infiltration: impaired energy metabolism accompanied by chronic inflammation and an altered tissue microenvironment. Notably, iron is a core component of key enzymes in the mitochondrial oxidative phosphorylation process, and its metabolic abnormalities can directly impact energy metabolism efficiency. Furthermore, chronic inflammation may also induce iron metabolic imbalance by regulating the expression of iron transport-related molecules.

Focused analysis of the regulatory patterns of iron metabolism-related proteins revealed that, in the total population, the protein levels of Transferrin (TF), Transferrin Receptor (TFRC), Ferritin Heavy Chain 1 (FTH1), Ferritin Light Chain (FTL), Cytochrome b Reductase 1 (CYBRD1), and Ceruloplasmin (CP) were significantly correlated with IMFF (**Fig. 1d**). This positive correlation was also observed in the male subgroup (**Fig. 1e**); in the female subgroup, the protein levels of TF, CYBRD1, and CP were similarly significantly positively correlated with IMFF (**Fig. 1f**). Statistical analysis confirmed that the levels of these iron metabolism-related proteins showed consistent trends across all groups (**Fig. 1g**). The widespread upregulation of iron metabolism-related proteins may represent a compensatory response of the body to systemic ID, aiming to enhance iron uptake and utilization, as exemplified by the upregulation of TFRC for iron uptake and CYBRD1 for iron transport.

In response to ID, cellular iron homeostasis is maintained through Nuclear Receptor Coactivator 4 (NCOA4)-mediated ferritinophagy. However, in the High group, ferritin (FTH1 and FTL) levels were significantly elevated. This paradoxical observation suggests that the ferritin degradation-iron release process may be impaired, leading to reduced levels of bioavailable iron and subsequent downregulation of iron utilization-related proteins. The Senescence-Associated Secretory Phenotype (SASP) in senescent cells can impair the ferroptosis process, which has been shown to impede NCOA4-mediated ferritinophagy.^[33,34]^ Consistent with this, our proteomic results also showed significant upregulation of chronic inflammation-related pathways, including complement and coagulation cascades. We further discovered that the protein level of NCOA4 was significantly downregulated in the skeletal muscle of the High group, accompanied by significant upregulation of FTH1 protein levels (**Supplementary Fig.5**). Given this, iron utilization in the skeletal muscle of elderly populations may be insufficient. Therefore, it is imperative to investigate the impact of iron insufficiency on skeletal muscle function. Under normal conditions, iron in the Labile Iron Pool (LIP) rapidly enters mitochondria for the synthesis of iron-containing proteins, while excess Fe²⁺ is either resequestered into ferritin or exported via ferroportin.^[35]^ However, measuring LIP levels only reflects the transient intracellular iron content and fails to accurately assess the cellular iron utilization status.^[36]^ Thus, analyzing the expression changes of iron utilization-related proteins is key to evaluating cellular iron demand. To determine the potential cause of the upregulation of iron metabolism-related proteins, we analyzed the correlation between iron utilization-related proteins and IMFF (**Fig. 1h, i, j**). These proteins included Myoglobin (MB), Aconitase 2 (ACO2), NADH Dehydrogenase (Ubiquinone) Iron-Sulfur Protein 1/2/3 (NDUFS1/2/3) and PHD2. The analysis revealed significant negative correlations between the levels of these iron utilization-related proteins and IMFF in the total population and in both male and female subgroups. Further analysis demonstrated that the levels of these proteins were significantly downregulated in the High group compared to the Low group across the total population and both genders (**Fig. 1k**). The significant downregulation of iron utilization-related proteins suggests severe impairment in the actual iron utilization process. Collectively, these findings indicate that skeletal muscle with high fat infiltration exhibits iron metabolism disorders, as evidenced by impaired ferritin degradation and compromised iron release, which ultimately presents as ID.

### 2.2. Reduced Muscle Function and Increased Fat Infiltration in ID Mice After Muscle Injury

To investigate the effects of ID on muscle function and fat infiltration, we first established a systemic iron homeostasis disruption model in mice via a low-iron diet as previously described.^[37]^ Subsequently, gastrocnemius muscle injury was induced by glycerol injection. This muscle was specifically selected due to its critical role in locomotion and strength, which are key indicators of skeletal muscle functional integrity,^[38]^ and its common use as a target site for inducing fat infiltration in mouse models.^[39]^ Glycerol was chosen for its well-characterized ability to induce muscle damage accompanied by fat infiltration, a hallmark of impaired muscle repair.^[40]^ Four weeks of low-iron diet feeding induced significant alterations in hematological parameters and iron-related indices in ID mice compared with iron-normal (IN) mice fed a standard diet. Specifically, ID mice exhibited a significant reduction in hemoglobin relative to IN mice. Concomitantly, marked decreases were observed in hematocrit, mean corpuscular volume, and mean corpuscular hemoglobin in ID mice (**Supplementary Fig.6a**), while red blood cell distribution width was significantly elevated (Supplementary Fig.6a). Beyond these hematological changes, ID mice also showed significant reductions in serum iron (**Supplementary Fig.6b**), muscle total iron (**Supplementary Fig.6c**), and muscle non-heme iron (**Supplementary Fig.6d**) compared with IN mice. Calcein fluorescence assays for the LIP, where higher fluorescence intensity indicates lower bioavailable iron,^[41]^ further confirmed intracellular iron depletion in ID mice (**Supplementary Fig.7a**). Specifically, mean fluorescence intensity was significantly elevated in endothelial cells (**Supplementary Fig.7b**), immune cells (**Supplementary Fig.7c**), SCs (**Supplementary Fig.7d**), and FAPs (**Supplementary Fig.7e**). Collectively, these results validated the successful establishment of the ID model.

To evaluate the impact of ID on post-injury skeletal muscle functional recovery, IN and ID mice were subjected to glycerol-induced muscle injury, with control mice receiving saline injection. All functional assessments and analyses were performed 4 weeks after modeling, as outlined in the experimental workflow (**Fig. 2a**). Behavioral tests confirmed functional deficits associated with ID during muscle recovery. CatWalk gait analysis was conducted to assess locomotor function; representative hindlimb footprint distributions and 3D pressure mapping (**Fig. 2b**) provided visual evidence of gait impairments. Quantitative analysis of gait parameters revealed that the ID injury group had significant reductions in maximum contact area (**Fig. 2c**), maximum intensity (**Fig. 2d**), and swing speed (**Fig. 2e**) compared with the IN injury group, indicating compromised locomotor coordination and hindlimb weight-bearing capacity. In addition, the ID injury group exhibited significantly lower maximal grip strength (**Fig. 2f**), reduced total treadmill exercise distance (**Fig. 2g**), and shorter rotarod retention time (**Fig. 2h**) relative to the IN injury group. Collectively, these quantitative functional assay results demonstrate that ID exacerbates functional impairments following muscle injury. H&E staining showed minimal fat deposition and evident repair in the IN injury group, whereas the ID injury group exhibited significantly increased fat accumulation in the injured area (**Fig. 2i**). Flow cytometry analysis indicated a significant increase in the FAP fraction and a significant decrease in the SC fraction in the ID injury group compared with the IN injury group, with no significant differences in other cell populations (**Fig. 2j, Supplementary Fig. 8a, 8b**). Taken together, these findings demonstrate that ID impairs functional recovery and exacerbates fat infiltration in injured muscle tissue. Given the critical role of FAPs in muscle fat infiltration and their significant upregulation in injured muscle from ID mice, this suggests that ID may increase fat infiltration by promoting the aberrant differentiation of FAPs.

### 2.3. ID Links HIF Pathway Activation in FAPs to Enhanced Skeletal Muscle Fat Infiltration

To investigate the functional alterations of FAPs under ID and their underlying mechanisms, FAPs were first isolated from the injured gastrocnemius muscles of IN and ID mice at 4 weeks post muscle injury induction (**Fig. 3a**), and flow cytometry verified the purity of these FAPs as 93.10% in the IN group and 83.40% in the ID group (**Supplementary Fig.9a**). Transcriptome sequencing of the purified FAPs revealed that the PCA plot (**Supplementary Fig.9b**) showed distinct separation of transcriptomic profiles between IN and ID groups. PC1 explained 50.09% of variance and PC2 accounted for 22.87%, with tight intra-group sample clustering. Meanwhile, the correlation heatmap (**Supplementary Fig.9c**) displayed high transcriptomic similarity among intra-group biological replicates (R²≥0.98), confirming excellent experimental reproducibility and reliable data quality. Additionally, the Venn diagram (**Supplementary Fig.9d**) illustrated that 11,233 genes (94.14%) were co-expressed in both groups, 386 genes (3.23%) were unique to IN FAPs, and 313 genes (2.62%) were specific to ID FAPs, verifying that ID induced distinct transcriptomic changes. Differentially expressed genes (DEGs) were screened using the criteria of |log₂(Fold Change)| ≥ 0.585 and p ≤ 0.05 (**Supplementary Fig.9e**), leading to the identification of 984 DEGs (456 upregulated and 528 downregulated in ID FAPs relative to IN FAPs), which provided a molecular basis for analyzing FAP functional changes under ID. KEGG pathway enrichment analysis (**Fig. 3b, 3c**) showed that upregulated DEGs in ID FAPs were primarily enriched in pathways including the HIF-1 signaling pathway and peroxisome proliferator-activated receptor (PPAR) signaling pathway, while downregulated DEGs were significantly enriched in Metabolic pathways and the Cell cycle signaling pathway. Finally, a heatmap of key functional genes (**Fig. 3d**) demonstrated distinct expression patterns in ID FAPs compared to IN FAPs: genes related to adipogenic differentiation (e.g., *Pparg*, *Adipoq*, *Cd36*), iron metabolism (e.g., *Hmox1*, *Fth1*), and HIF-1 signaling (e.g., *Egln1*, *Egln3*) were upregulated, while cell proliferation-related genes (e.g., *Ccnb1*, *Cdk1*, *Bub1b*) were downregulated, implying potential links between ID, HIF-1α activation, and FAP adipogenic differentiation.

To confirm the critical role of HIF-1α in muscular fat infiltration, we further analyzed the changes in HIF-1α protein levels in skeletal muscle samples with different degrees of fat infiltration. Western blot analysis (**Fig. 3e**) confirmed that the protein levels of HIF-1α was significantly upregulated in the High group. Immunofluorescence staining targeting HIF-1α and PDGFRα demonstrated stronger HIF-1α fluorescence, more dispersed PDGFRα⁺ FAPs, and a higher number of HIF-1α⁺/PDGFRα⁺ co-localized cells in the High group compared to the Low group (**Fig. 3f**). Quantitative analysis validated that the High group exhibited significantly increased abundance of PDGFRα⁺ FAPs (**Fig. 3g**), and a higher percentage of HIF-1α⁺ cells among PDGFRα⁺ FAPs (**Fig. 3h**), indicating FAP-specific HIF-1α activation in skeletal muscle with high fat infiltration.

### 2.4. Upregulation of HIF-1α Mediates ID-Induced Enhanced Adipogenic Differentiation of FAPs

To simulate ID in vitro and validate the regulatory effects of iron depletion on FAP function, FAPs were treated with deferoxamine (DFO) at concentrations of 0, 1, 2, 5, 10, and 20 μM, and cell viability was assessed to ensure the safety of the selected concentrations (**Supplementary Fig.10a, S10b**). Calcein and FerroOrange fluorescence assays for LIP and intracellular Fe²⁺ levels, respectively (**Supplementary Fig.10c-f**), showed that DFO treatment significantly increased Calcein mean fluorescence intensity (**Supplementary Fig.10c, S10d**) and decreased FerroOrange MFI (**Supplementary Fig.10e, S10f**) in a concentration-dependent manner, validating effective induction of intracellular iron depletion in FAPs. Based on this iron depletion model, quantitative real-time reverse transcription polymerase chain reaction (qRT-PCR) analysis (**Fig. 4a–f**) demonstrated that as DFO concentration increased, the mRNA levels of adipogenic differentiation markers (*Pparg*, Fig. 4a; *Adipoq*, Fig. 4b) and the iron metabolism-related gene (*Hmox1*, Fig. 4c) were significantly upregulated, while the mRNA levels of cell proliferation-related genes (*Ccnb1*, Fig. 4d; *Cdk1*, Fig. 4e; *Bub1b*, Fig. 4f) were significantly downregulated in a concentration-dependent manner. These results indicate that iron depletion promotes adipogenic differentiation of FAPs and inhibits their proliferation. Western blot analysis of FAPs treated with 0, 1, and 2 μM DFO (**Fig. 4g**) revealed significant reductions in FTH1 protein levels (**Fig. 4h**), alongside marked increases in PPARγ protein levels (**Fig. 4i**) and HIF-1α protein levels (**Fig. 4j**). These results align with transcriptomic changes and confirm that iron depletion regulates HIF-1α activation and the expression of adipogenic-related proteins. Finally, Nile red staining (**Fig. 4k**) and quantitative analysis of staining intensity (**Fig. 4l**) revealed that low concentrations of DFO promoted lipid droplet formation in FAPs, while high concentrations (>10 μM) exerted an inhibitory effect. This bidirectional regulation may be associated with reduced cell viability induced by severe iron depletion under high DFO concentrations. Collectively, these findings indicate that ID alters the transcriptomic profile of FAPs, and in vitro iron depletion regulates FAP adipogenic differentiation in a concentration-dependent manner where mild iron depletion promotes adipogenesis while severe iron depletion inhibits it, while simultaneously suppressing proliferation. This process is closely associated with activation of the HIF-1α signaling pathway, which may be a key mechanism underlying increased fat infiltration in injured muscles of ID mice.

**Fig. 4.**
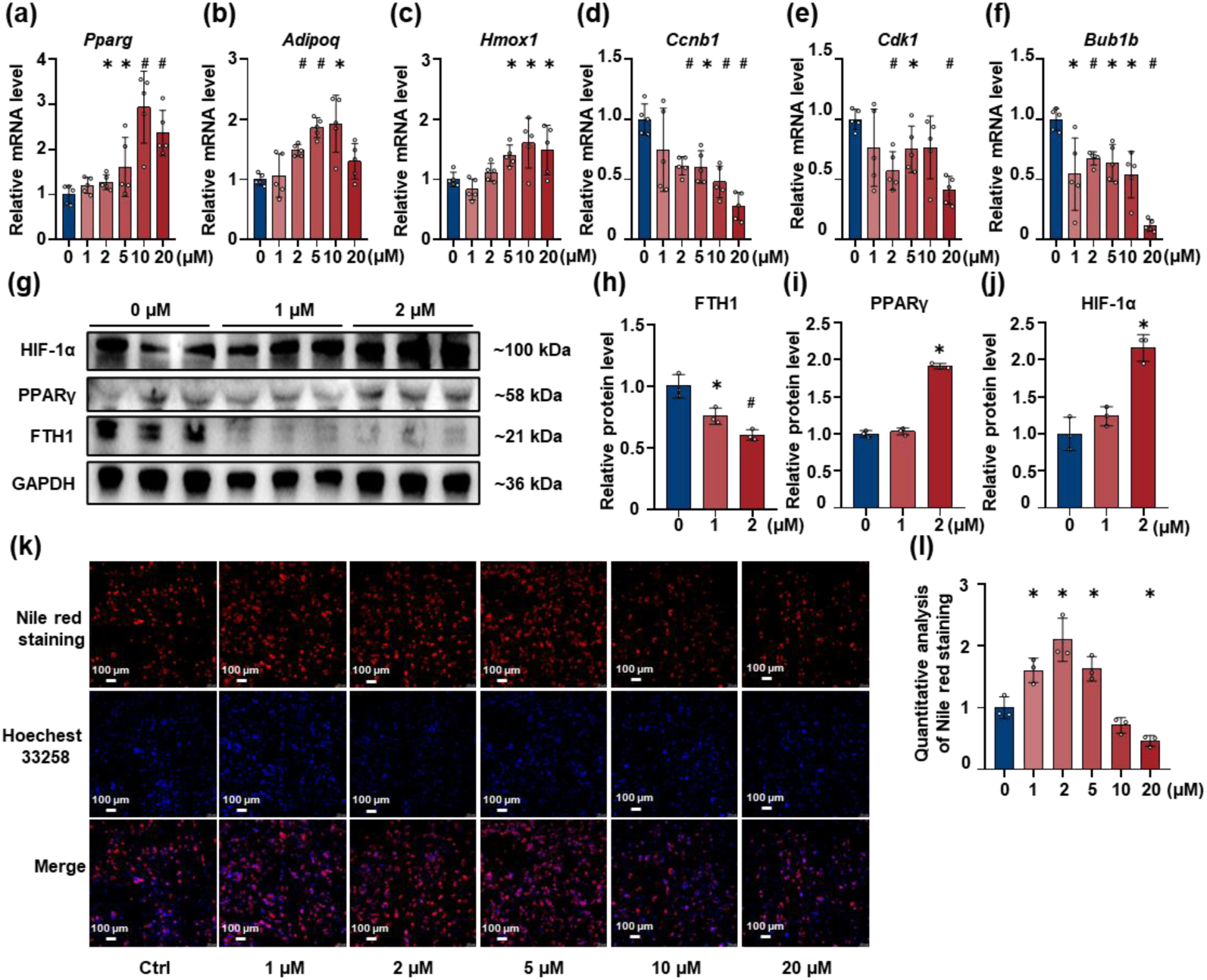
Functional analysis of FAPs treated with gradient concentrations of DFO in vitro. (a–f) Relative mRNA levels of adipogenic (a: *Pparg*, b: *Adipoq*), iron metabolism (c: *Hmox1*), and cell proliferation (d: *Ccnb1*, e: *Cdk1*, f: *Bub1b)* genes in FAPs treated with different concentrations of DFO (0, 1, 2, 5, 10, 20 μM), detected by qRT-PCR (n=5). (g) Western blot analysis of HIF-1α, PPARγ, and FTH1 protein levels in FAPs treated with 0, 1, and 2 μM DFO, with GAPDH as a loading control. (h–j) Quantitative analysis of the relative protein levels of FTH1 (h), PPARγ (i), and HIF-1α (j) based on the Western blot results in (j) (n=3). (k) Nile red staining (red, lipid droplets), Hoechst 33258 staining (blue, nuclei), and merged images of FAPs treated with different DFO concentrations (0, 1, 2, 5, 10, 20 μM). (l) Quantitative analysis of Nile red staining intensity corresponding to panel (k) (n=3). * indicates a significant difference (p < 0.05) compared to the 0 μM DFO group; # indicates a significant difference (p < 0.001) compared to the 0 μM DFO group.

### 2.5. Inhibition of HIF-1α Reverses Iron Depletion-Enhanced FAP Adipogenic Capacity

To verify the role of HIF-1α in mediating iron depletion-induced adipogenic differentiation of FAPs, we used PX478, a specific HIF-1α inhibitor; after assessing FAP viability following treatment with varying concentrations of PX478, 2 μM was selected for subsequent experiments (**Supplementary Fig.11a, S11b**). To verify whether HIF-1α activation mediates iron depletion-induced FAP adipogenic differentiation, FAPs were divided into three groups: the control group (Ctrl), the iron depletion group (DFO-treated), and the iron depletion combined with HIF-1α inhibition group (DFO and PX478 co-treated). qRT-PCR analysis (**Fig. 5a–f**) showed that relative to the Ctrl group, the DFO group had significantly higher mRNA levels of adipogenic markers *Pparg* **(Fig. 5a**) and *Adipoq* (**Fig. 5b**), as well as the iron metabolism-related gene *Hmox1* (**Fig. 5c**), and significantly lower mRNA levels of cell proliferation/cycle-related genes *Ccnb1* (**Fig. 5d**), *Cdk1* (**Fig. 5e**), and *Bub1b* (**Fig. 5f**). Notably, these expression trends were significantly reversed in the DFO and PX478 co-treated group compared with the DFO group, with reduced *Pparg* and *Adipoq* mRNA levels and increased *Ccnb1*, *Cdk1*, and *Bub1b* mRNA levels, suggesting that HIF-1α inhibition counteracts the regulatory effects of iron depletion on FAP gene expression. Western blot analysis (**Fig. 5g**) of HIF-1α, NCOA4, PPARγ, and FTH1 protein levels. Relative to the Ctrl group, the DFO group exhibited marked upregulation of HIF-1α and PPARγ, alongside the ID-typical elevation of NCOA4 and reduction of FTH1. Conversely, the DFO+PX478 co-treatment group had drastically lower HIF-1α and PPARγ levels than the DFO group, while NCOA4 and FTH1 levels remained comparable to those in the DFO group. These results confirm PX478’s effective inhibition of HIF-1α activation, and more importantly, they show that HIF-1α suppression under ID directly reverses FAP adipogenic capacity. To assess this effect on FAP adipogenic differentiation, Nile red staining revealed far more lipid droplets in the DFO group than in the Ctrl group. By contrast, DFO+PX478 co-treatment markedly reduced lipid droplet number and size (**Fig. 5h**). Quantification of Nile red fluorescence intensity validated this pattern. Intensity was significantly higher in the DFO group than in the Ctrl group, but it was drastically lower and approached Ctrl levels in the DFO+PX478 group relative to the DFO group (**Fig. 5i**). Collectively, these data establish HIF-1α activation as a key mediator of iron depletion-induced FAP adipogenic differentiation. PX478-mediated HIF-1α inhibition reverses iron depletion-enhanced FAP adipogenic capacity, normalizes aberrant expression of molecules related to adipogenesis and cell proliferation, and reduces FAP lipid droplet accumulation. This confirms the HIF-1α pathway’s central role in regulating FAP function under ID.

**Fig. 5.**
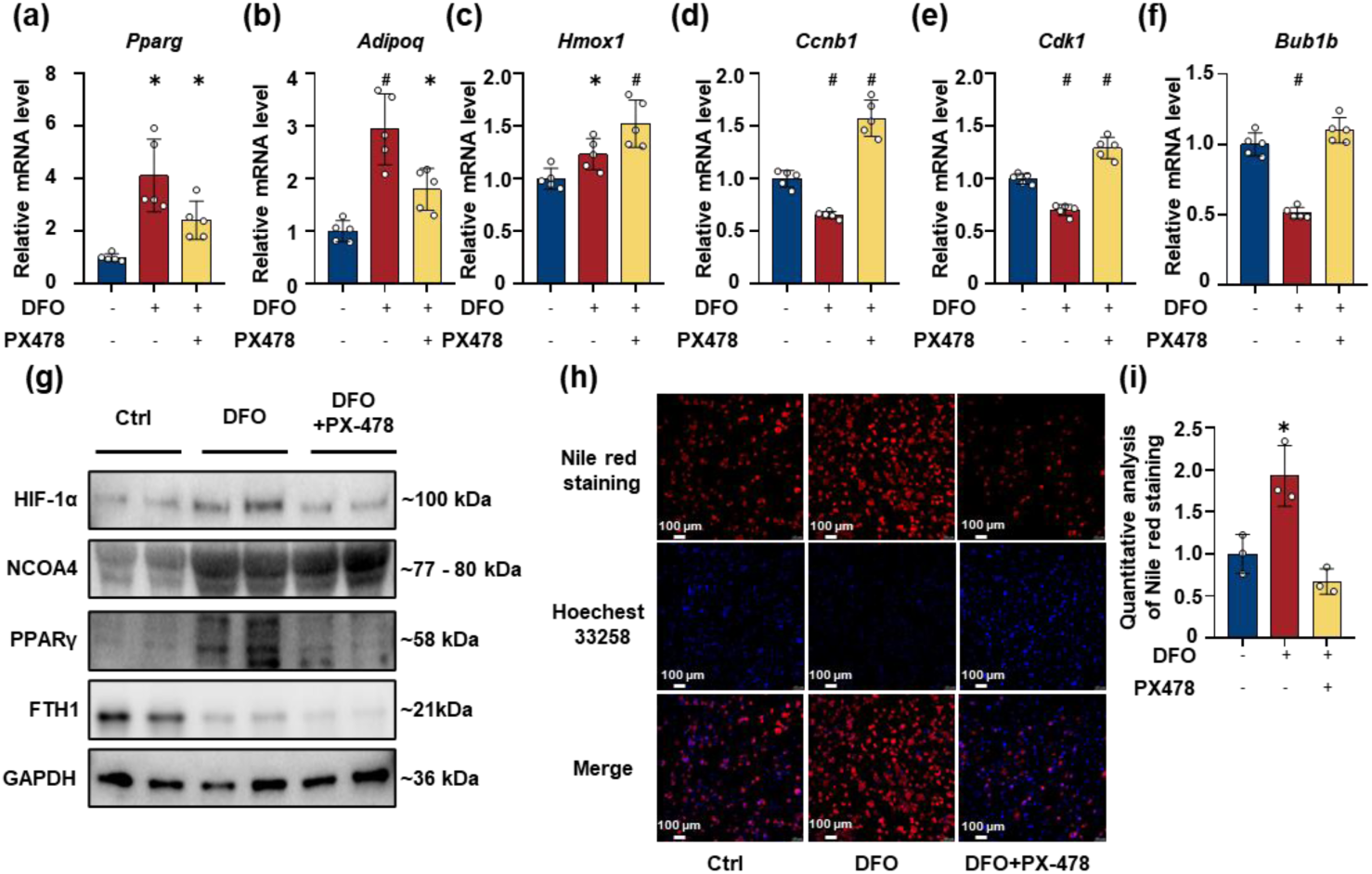
In vitro functional validation of FAPs treated with DFO and HIF-1α inhibitor PX478. (a–f) Relative mRNA levels of adipogenic (a: *Pparg*, b: *Adipoq*), iron metabolism-related (c: *Hmox1*), and cell proliferation/cycle-related (d: *Ccnb1*, e: *Cdk1*, f: *Bub1b*) genes in FAPs from the Ctrl group, DFO-treated group, and DFO and PX478 co-treated group, detected by qRT-PCR. (g) Western blot analysis of HIF-1α, NCOA4, PPARγ, and FTH1 protein levels in FAPs from the Ctrl, DFO-treated, and DFO and PX478 co-treated groups, with GAPDH used as a loading control. (h) Nile red staining (red, lipid droplets), Hoechst 33258 staining (blue, nuclei), and merged images of FAPs from the Ctrl, DFO-treated, and DFO and PX478 co-treated groups. Scale bar = 100 μm. (i) Quantitative analysis of Nile red staining intensity corresponding to panel (n=3). * indicates a significant difference (p < 0.05) compared to the Ctrl group; # indicates a significant difference (p < 0.001) compared to the Ctrl group.

### 2.6. HIF-1α Inhibition by PX478 Improves Muscle Function Recovery in ID Mice After Glycerol-Induced Injury

To confirm whether targeting HIF-1α with PX478 can reverse impaired muscle repair and abnormal iron metabolism induced by ID post-injury, IN and ID mice were each divided into three subgroups: control (Ctrl, saline-injected), injury (glycerol-induced muscle injury), and PX478-treated (injury treated with the HIF-1α inhibitor PX478), with analyses of behavioral function, histological morphology, key protein expression, and FAP/SC proportions performed (**Fig. 6a**). Gait analysis (**Fig. 6b–e**) confirmed functional improvement, as the ID-Injury group had significantly reduced max contact area, mean intensity, and swing speed vs. IN-Injury, while these parameters were significantly recovered in the ID-PX478 group vs. ID-Injury. In addition, the ID-Injury group had significantly lower maximal grip strength (**Fig. 6f**) than both the IN-Injury and ID-Ctrl groups, with PX478 treatment reversing this deficit to approach the IN-Injury group level. Similar trends were observed in treadmill test results **(Fig. 6g**) and rotarod test results (**Fig. 6h**), revealing that the ID-Injury group exhibited a significant reduction in treadmill movement distance and rotarod residence time compared with the IN-Injury group, and these impairments were rescued by PX478 treatment. H&E staining (**Fig. 6i**) showed no fat deposition in the IN-Ctrl group, minimal fat infiltration in the IN-Injury group, and prominent fat deposition in the ID-Injury group; notably, the ID-PX478 group exhibited reduced fat deposition compared with the ID-Injury group, while the IN-PX478 group showed a similar degree of fat infiltration to the IN-Injury group, suggesting that HIF-1α inhibition mitigates fat infiltration in injured ID muscles. Western blot analysis (**Fig. 6j**) of HIF-1 signaling, iron metabolism, and adipogenesis-related proteins demonstrated that the ID-Injury group had significantly increased HIF-1α and PPARγ levels alongside decreased NCOA4 and FTH1 levels compared with the IN-Injury group, while PX478 treatment significantly reversed these abnormalities in the ID-PX478 group with no significant changes observed in the IN-PX478 group relative to the IN-Injury group, confirming that PX478 specifically targets HIF-1α-mediated dysregulation in ID mice. Flow cytometry analysis of FAP and SC proportions (**Supplementary Fig.12a–c**) revealed that the FAP fraction in the ID-Injury group was higher than that in the IN-Injury group. Notably, PX478 treatment significantly reduced the FAP fraction and increased the SC fraction, thereby correcting this cellular proportion imbalance. This finding indicates that PX478 restores the balance of functional cell populations, which is critical for muscle repair. Collectively, these results demonstrate that the HIF-1α inhibitor PX478 exerts multiple protective effects in ID mice after muscle injury, including rescue of impaired muscle function, alleviation of fat deposition, normalization of HIF-1α-mediated adipogenesis-related protein expression, and reduction in FAP number, confirming HIF-1α in FAPs as a key therapeutic target for improving muscle repair under ID conditions.

**Fig. 6.**
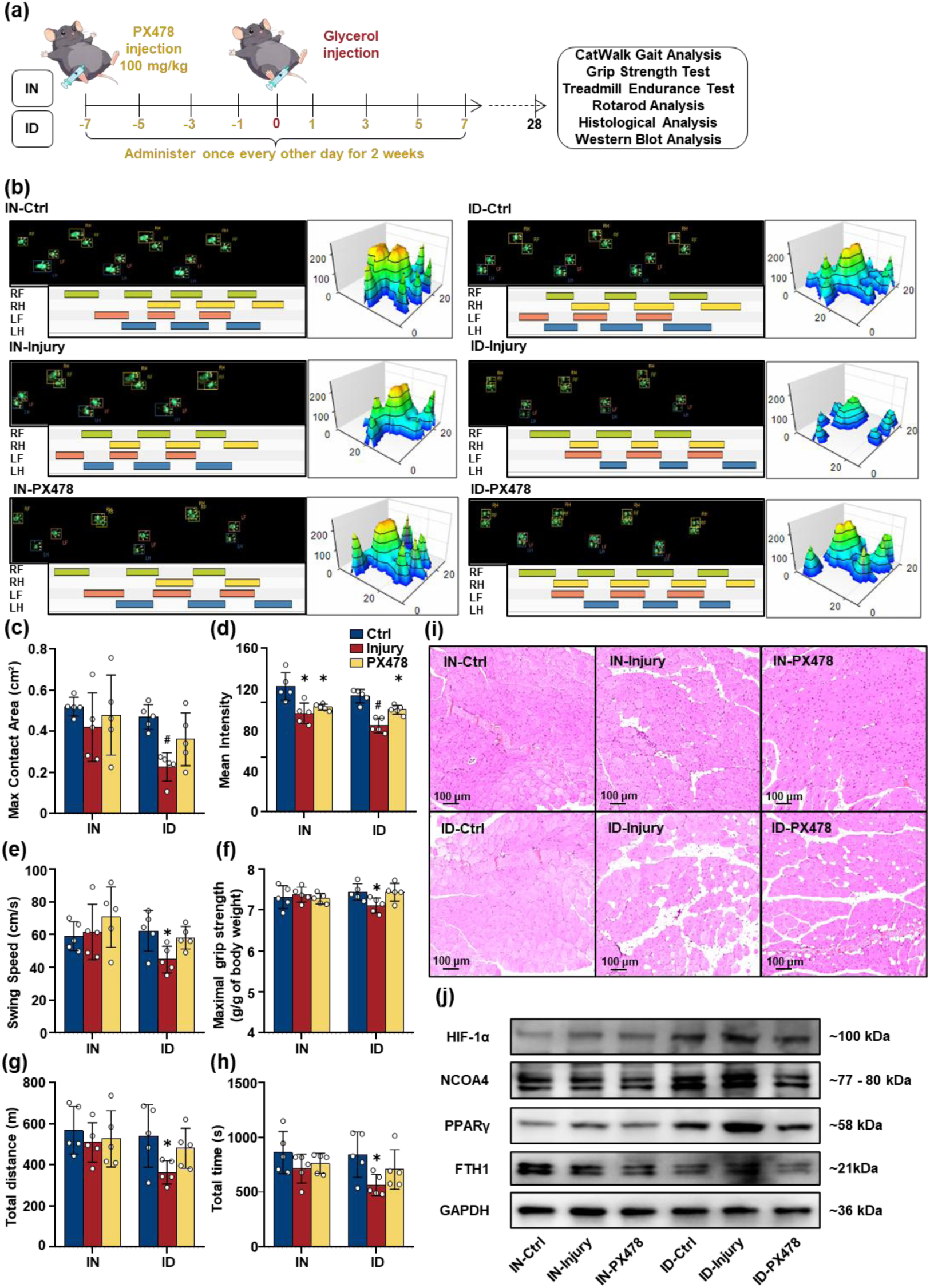
Muscle function and morphological analysis of muscle recovery after glycerol-induced injury and HIF-1α inhibition by PX478 in IN and ID models. (a) Experimental timeline: IN and ID mice were administered PX478 (100 mg/kg, once every other day for 2 weeks, initiated 7 days prior to injury). Muscle injury was induced via intramuscular injection of 100 μL 50% (v/v) glycerol at day 0. At 28 days post-injury, analyses including CatWalk gait assessment, grip strength test, treadmill endurance test, rotarod test, histological examination, and Western blot were conducted. (b) CatWalk gait analysis visualization: Representative hindlimb footprint distributions (top panels), strip plots of gait parameters (bottom panels), and 3D pressure distribution maps of the left hindlimb (right panels) for IN and ID mice across three groups: control (Ctrl, no injury), glycerol-induced injury (Injury), and injury + PX478 treatment (PX478). (c–e) Gait analysis results: (c) Max contact area (cm²), (d) Mean intensity, and (e) Swing speed (cm/s) in Ctrl, Injury, and PX478 groups of IN and ID models (n=5). (f) Maximal grip strength (g/g body weight) in control (Ctrl), injury (glycerol-injected), and PX478-treated groups of IN and ID models (n=5). (g) Treadmill Test: Total Movement Distance (m) in the Control (Ctrl), Injury, and PX478-Treated Groups of IN and ID Mouse Models (n=5). (h) Rotarod Test: Residence Time (s) in the Control (Ctrl), Injury, and PX478-Treated Groups of IN and ID Mouse Models (n=5). (i) H&E staining of muscle tissues from IN-Ctrl, IN-Injury, IN-PX478, ID-Ctrl, ID-Injury, and ID-PX478 groups. Scale bar = 100 μm. (j) Western blot analysis of HIF-1α, NCOA4, PPARγ, and FTH1 protein levels in IN-Ctrl, IN-Injury, IN-PX478, ID-Injury, and ID-PX478 groups, with GAPDH as a loading control. * and # indicate significant differences (p < 0.05, p < 0.001) between specified groups.

## 3. Discussion

In this study, we demonstrate that ID drives skeletal muscle fat infiltration via HIF-1α-mediated aberrant FAP differentiation. Our findings identify a novel functional axis where ID stabilizes HIF-1α to direct FAP adipogenic differentiation and subsequent muscle steatosis. While these results advance our understanding of ID-linked muscle fat infiltration, several key aspects merit further discussion.

First, the mechanisms underlying functional ID and iron sequestration in muscle warrant further mechanistic investigation. While our data indicate that NCOA4 protein downregulation in skeletal muscle with high fat infiltration may elevate intramuscular ferritin levels, thereby reducing bioavailable iron pools and decreasing the expression of iron-utilizing proteins to ultimately elicit a functional iron-deficient phenotype, this represents only one contributing factor. Notably, chronic inflammation, a common comorbidity of ID and aging, exerts a pivotal role by upregulating hepcidin expression;^[42]^ this hormone blocks iron export from macrophages and potentially muscle cells alike.^[43,44]^ Such inflammatory iron sequestration may further exacerbate local ID despite overt tissue iron accumulation.^[45,46]^ Future investigations focusing on the crosstalk between inflammatory cytokines (e.g., IL-6) and iron regulatory proteins (e.g., ferroportin) in ID-affected muscle will be essential to fully unravel this paradoxical iron metabolic phenotype.

Second, the dual role of HIF-1α in muscle homeostasis necessitates a more nuanced interpretation. While our study implicates HIF-1α as a key driver of FAP adipogenic differentiation under ID conditions, it is essential to acknowledge its context-dependent protective functions. Acute HIF-1α activation promotes glycolytic metabolism and cell survival during hypoxia or injury.^[21–24]^ However, in various chronic diseases, long-term stabilization of HIF-1α may contribute to disease pathogenesis. In neutrophils from patients with glycogen storage disease type Ib (GSD-Ib), increased HIF-1α expression acts as an upstream activator to promote PPAR-γ expression.^[28]^ Additionally, alcohol feeding can activate hepatic HIF-1α, thereby facilitating lipid accumulation.^[29]^ In non-alcoholic steatohepatitis, HIF-1α levels are elevated in hepatic macrophages and circulating monocytes of patients; this further promotes inflammation via activation of the NF-κB pathway, which in turn impairs lipid metabolism.^[47]^ Furthermore, as a key transcription factor for dhrs3a, HIF-1α enhances dhrs3a expression, leading to increased retinol production catalyzed by Dhrs3. This activates the PPAR-γ pathway, alters lipid composition, and indirectly positively regulates PPAR-γ, thereby participating in the process of lipid droplet accumulation.^[27]^ In our study, long-term ID in skeletal muscle similarly induces HIF-1α accumulation, which triggers aberrant FAP differentiation and subsequent muscle fat infiltration. Understanding the temporal and spatial dynamics of HIF-1α activity in muscle pathologies may identify therapeutic windows for targeted intervention without compromising its physiological functions.

Third, the crosstalk between FAPs and SCs is a pivotal determinant of muscle regeneration and warrants expanded discussion. Our results show that ID expands FAP populations while reducing SC numbers and impairing their myogenic differentiation. FAPs’ autonomous differentiation into adipocytes is a well-established contributor to intramuscular fat accumulation, and emerging evidence from skeletal muscle research underscores that FAPs modulate SC behavior primarily through paracrine signaling.^[17]^ Consistent with the documented regulatory roles of FAP-secreted factors in muscle homeostasis, we speculate that ID-skewed FAPs may secrete inhibitory molecules including TGF-β and WNT pathway inhibitors, which impede SC activation and self-renewal or even skew their fate toward non-myogenic lineages. Elucidating the precise paracrine communication mechanisms underlying this FAP-SC crosstalk could uncover novel therapeutic targets for preserving muscle regenerative capacity under ID conditions.

Finally, the therapeutic potential of HIF-1α inhibition (PX478) must be balanced with safety concerns. While PX-478 effectively reversed muscle steatosis in our models, systemic HIF-1α blockade risks impairing erythropoiesis, angiogenesis, and wound healing.^[21–24]^ Therefore, alternative strategies merit exploration: (1) muscle-specific HIF-1α modulation to avoid systemic toxicity; (2) restoring iron utilization via NCOA4 agonists to alleviate functional ID without suppressing HIF-1α globally; or (3) targeted iron delivery systems (e.g., nanoparticle-based iron supplementation) to bypass impaired iron export pathways. Preclinical studies comparing these approaches are urgently needed to inform clinical translation.

In summary, our study uncovers a HIF-1α-dependent mechanism linking ID to muscle fat infiltration, yet several critical knowledge gaps remain to be filled. Future investigations that focus specifically on the inflammatory drivers of iron sequestration, the temporal dynamics of HIF-1α, cell-cell interactions and safer therapeutic strategies will help address these limitations, enabling us to more effectively translate these findings into clinical interventions for ID-related muscle disorders.

## 4. Conclusion

This study, integrating clinical sample analysis and an ID model in aged mice, systematically delineates the pivotal role of ID in skeletal muscle fat infiltration and impaired regeneration. Consistent clinical and experimental evidence from the aged mouse ID model converges to confirm that ID-mediated stabilization of HIF-1α expression is a key trigger for the activation of the ID-HIF-1α-FAPs signaling axis. This axis drives aberrant adipogenic differentiation of FAPs, disrupts the critical balance between FAPs and SCs, and ultimately culminates in intramuscular lipid accumulation and functional deterioration. Pharmacological inhibition of HIF-1α using PX478 effectively reversed ID-induced pathological adipogenesis and restored muscle regenerative capacity, validating this axis as a promising therapeutic target. By incorporating clinical evidence with the aged mouse ID model, our study not only elucidates the mechanisms underlying ID-associated muscle pathologies but also provides novel insights into interventions for elderly patients and those with iron dysmetabolism.

## 5. Experimental Methods

### Study Design and Ethical Approval

A single-center cross-sectional study was conducted at Beijing Jishuitan Hospital. This study stringently complied with the principles outlined in the Declaration of Helsinki and was approved by the Institutional Ethics Committee of Beijing Jishuitan Hospital (Ethical approval number: K2023-138-00). Written informed consent was obtained from all participants prior to study enrollment.

### Participants

Eligible participants were those who met the following requirements: aged between 60 and 90 years, with a primary diagnosis of lumbar degenerative disease, and voluntary participation accompanied by signed informed consent, while individuals were excluded if they satisfied any of the following conditions: body mass index (BMI) ≤ 18 kg/m² or ≥ 30 kg/m², severe pain during spinal movement, Oswestry Disability Index (ODI) > 80%, severe spinal deformity (congenital scoliosis with a Cobb angle > 10° or lumbar spondylolisthesis ≥ Grade 1), a history of lumbar trauma, spinal surgery, or infectious diseases, a history of neurological, neuromuscular, autoimmune, tumor, or psychiatric disorders, or ongoing use of medications impacting muscle structure or function. A total of 781 patients were initially recruited and following stringent screening against the previously outlined inclusion and exclusion criteria, 330 patients were qualified for further assessment; ultimately, 174 participants who underwent comprehensive imaging and muscle strength data collection were included in the final analysis.

### Clinical Data and Functional Assessments

Demographic data incorporating age, gender, and BMI were collected from all participants, with spinal function evaluated by the ODI and the severity of low back pain gauged using the Visual Analog Scale (VAS), while sarcopenia was thoroughly evaluated via a battery of tests encompassing the SARC-F scale, Short Physical Performance Battery (SPPB), gait speed measurement, grip strength testing, and chair stand test. Grip strength was measured quantitatively using a Jamar handheld dynamometer.^[48,49]^ In accordance with the 2019 criteria of the Asian Working Group for Sarcopenia (AWGS 2019), possible sarcopenia was defined as low grip strength—males have a grip strength of < 28 kg and females < 10 kg—coupled with compromised physical performance, which refers to a gait speed of < 1.0 m/s, an SPPB score of ≤ 9, or a chair stand test duration of ≥ 12 s.^[50]^ And ultimately, 84 male and 90 female participants who furnished comprehensive imaging and muscle strength data sets were included for the final analysis, with the basic demographic and clinical features of these 174 participants depicted in Supplementary Fig.1a-1b.

### Imaging Data Acquisition and Analysis

For MRI scanning, lumbar T2-weighted images were acquired using a 3.0 T MRI scanner (Ingenia 3.0 T, Philips), with axial images corresponding to the L4/5 intervertebral disc level chosen for further analytical procedures, while for the quantification of fat and muscle components, the Otsu thresholding algorithm was employed to automatically differentiate between fat and water signals within the paraspinal muscles on MRI images, thereby isolating intramyocellular fat (IMF) and perimuscular fat (PMF); the IMFF was calculated as IMFF = [IMF / (functional cross-sectional area [fCSA] + IMF)] × 100%, and the perimuscular fat fraction (PMFF) was computed as PMFF = (PMF / total cross-sectional area [TCSA]) × 100%, with muscle mass expressed as height-adjusted muscle area including the PSMI (PSMI = paraspinal muscle fCSA / height², cm²/m²) and skeletal muscle index (SMI = cross-sectional skeletal muscle area / height², cm²/m²), and two spinal surgeons possessing 5 years of clinical practice and MRI reading expertise evaluated the images in an independent manner, with intraclass correlation coefficients (ICC) utilized to assess inter-observer and intra-observer consistency.

### Isokinetic Muscle Strength Testing

Prior to the commencement of testing, participants completed a 5-min walking session and a 2-min warm-up routine. Isokinetic movements, encompassing rotation, flexion, extension, and lateral flexion, were executed using a BioniX Sim3 Pro device (Physiomed, Germany) at an angular velocity of 15 °·s⁻¹. Each movement was performed for five consecutive repetitions with an inter-repetition interval of 20 s. Peak torque values were logged, and normalized muscle strength (peak torque/body weight, Nm/kg) was derived, including extension strength, rotational strength (mean value of the left and right sides), lateral flexion strength (mean value of the left and right sides), and average paraspinal muscle strength.

### Peripheral Blood Samples

Fasting peripheral blood samples (5 mL) were collected from each participant before surgery. Samples were centrifuged at 3000 × g for 10 min at 4 °C to separate serum, which was stored at -80 °C for subsequent serum iron detection. Whole blood samples were used for routine blood tests within 2 h of collection.

### Skeletal Muscle Samples

Paraspinal multifidus muscle biopsies were obtained during spinal surgery at the L4/5 intervertebral disc level. Fresh muscle tissue was immediately cleaned with sterile wet gauze to remove necrotic tissue, epimuscular fat, and excess tendons. Tissue samples were divided into 100 mg aliquots: one aliquot was fixed in 4% paraformaldehyde for histological analysis, and the remaining aliquots were snap-frozen in liquid nitrogen and stored at -80 °C for proteomic sequencing, tissue iron detection, and Western blot analysis.

### Proteomic Sequencing and Bioinformatics Analysis

Based on the previously acquired imaging findings (for fat infiltration quantification) and muscle strength measurement data, we selected samples categorized into two cohorts for proteomic sequencing: the high fat infiltration group (with fat infiltration degree > 15%, a threshold corresponding to significant reduction in skeletal muscle function) and the low fat infiltration group (with fat infiltration degree < 5%). A total of 79 samples were included in this analysis: 51 samples were assigned to the low fat infiltration group, while 28 samples belonged to the high fat infiltration group. Stratified by gender, this cohort comprised 46 male samples (35 allocated to the low group and 11 to the high group) and 33 female samples (16 in the low group and 17 in the high group). These selected samples were subsequently subjected to proteomic sequencing procedures detailed below.

For protein extraction and digestion, total protein was isolated from human tissue specimens using lysis buffer (Omicsolution, China). Roughly half of each sample was mixed with 300 μL buffer, mechanically disrupted with steel beads (35 Hz, 2 min × 2) and sonicated in an ice-water bath for 20 min. The homogenate was centrifuged (12000 rpm, 4 °C, 10 min) to collect supernatant, with protein concentration quantified via the BCA method (Enhanced BCA Kit, Beyotime Biotechnology, China). Thirty micrograms of protein was mixed with 20 μL reduction/alkylation reagent (Omicsolution, China) and incubated at 95 °C for 5 min at 1000 rpm. After cooling to room temperature, 15 μL protease reagent (Omicsolution, China) was added for digestion at 37 °C for 2 h at 1000 rpm, and digestion was terminated with 55 μL reagent (Omicsolution, China). Peptides were desalted using C18 cartridges (Omicsolution, China), concentrated via vacuum freeze centrifugation, and reconstituted in iRT-containing mobile phase A (Biognosys, CH) for quantification. For LC-MS/MS analysis, reconstituted peptides were separated by nano-UPLC (Evosep one, Denmark) coupled to a timsTOF Pro2 mass spectrometer (Bruker, Germany; nano-electrospray ion source) on a PePSep C18 column (1.9 μm, 150 μm × 15 cm, Bruker, Germany). Mobile phases were 0.1% formic acid in water (A) and acetonitrile (B). The column was equilibrated with 100% A, and peptides separated via gradient elution. Mass spectrometry used DIA-PASEF mode (100–1700 m/z scan range), with collision energy increasing linearly (20–59 eV) with ion mobility (1/K0: 0.6–1.6 Vs/cm²). For data processing, raw MS files were analyzed via Spectronaut v19.0.240604.62635 (Biognosys AG) against the uniprot_iRT_Homo sapiens_9606_reviewed_2024.fasta database. Parameters included trypsin (≤2 missed cleavages), fixed carbamidomethylation (C), variable acetyl (N-terminus) and oxidation (M), 20 ppm mass tolerance, and FDR ≤ 0.01 (peptide/PSM levels). Differentially abundant proteins were identified with |log₂FoldChange| > 0.585 and P-value < 0.05. KEGG pathway enrichment analysis was performed to explore biological processes.

### Animals and Experimental Design

Male C57BL/6N mice (18-month-old) were purchased from Beijing Vital River Laboratory Animal Technology Co., Ltd. Mice were randomly divided into two groups: the IN group fed a standard diet (iron content: 105.26 ppm) and the ID group fed a low-iron diet (iron content: 5.93 ppm) for 4 wk to establish the ID model, as previously described.^[37]^ After model establishment, both groups were further divided into three subgroups: the control (Ctrl, saline injection), injury (glycerol-induced muscle injury), and PX478-treated (injury + PX478 administration) groups. For muscle injury induction, 100 μL of 50% (v/v) glycerol solution was intramuscularly injected into the gastrocnemius muscle. For the PX478-treated group, PX478 (100 mg/kg body weight) was intraperitoneally injected every other day starting from day -7, with the treatment lasting for 2 wk. All animal care and experimental protocols were approved by the Animal Ethics Committee of Beijing Jishuitan Hospital, Capital Medical University (Ethical approval number: JSTLAEC2025-03-S16) and conducted in accordance with the Guide for the Care and Use of Laboratory Animals.

### Hematological and Iron Parameter Analysis

Blood samples were collected from mice via retro-orbital venous plexus puncture. For routine blood testing, 20 μL of the collected blood was diluted at a 1:100 ratio with a standard dilution buffer, and a complete blood count (CBC) analysis was subsequently performed using a hematology analyzer (Nihon Kohden Corporation, Tokyo, Japan). For the serum iron assay, the remaining volume of the collected blood was centrifuged at 3500 × g for 10 min to isolate serum, and serum iron concentration was measured using a Serum Iron Assay Kit (Nanjing Jiancheng Bioengineering Institute, China) in accordance with the manufacturer’s instructions. Briefly, blank, standard, and test tubes were prepared, with each tube containing 50 μL of distilled water, iron standard solution, or the isolated serum, respectively. After adding 150 μL of freshly prepared iron chromogenic reagent to each tube, the tubes were incubated in a boiling water bath for 5 min. Following centrifugation, the absorbance of the supernatants was measured at 520 nm using an ALLSHENG Feyond-A300 Microplate Reader (ALLSHENG, China).

### Non-heme iron detection

Reagents included: Chromagen stock solution (100 mg bathophenanthrolinedisulfonic acid disodium salt hydrate dissolved in 60 mL MilliQ water, mixed with 1.429 mL 70% thioglycolic acid and diluted to 100 mL); freshly prepared Chromagen working solution (1:5:5 mix of stock solution, saturated sodium acetate, MilliQ water); mixed acid (49.6 mL 36–38% hydrochloric acid, 20 mL 100% trichloroacetic acid diluted to 200 mL with MilliQ water). For detection, weighed tissue was placed in centrifuge tubes with 1 mL mixed acid, sealed, and incubated at 65 °C in a water bath for 24 h. After adding pre-soaked zirconia beads, homogenizing, and another 4 h water bath, samples were cooled, diluted to 1.5 mL with mixed acid, and centrifuged at 12000 rpm for 5 min. Tissue lysate (sample), mixed acid (blank), and iron standard solution (standard) were added to tubes with mixed acid and Chromagen working solution, incubated 10 min at room temperature. Absorbance of 100 μL samples (assayed in a 96-well plate) was measured at 535 nm using a microplate reader, and the iron content (Fe, expressed in μg/g wet tissue) was calculated as Fe

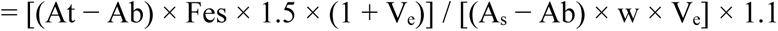

(Aₜ: sample OD; Ab: blank OD; Aₛ: average iron standard OD; Fes = 0.2492; w: tissue wet weight; Vₑ = 0.5).

### Total Iron Detection

Mouse muscle tissue samples were collected and weighed. The iron content was determined using a commercial kit (Elabscience Biotechnology Co., Ltd., China) according to the manufacturer’s instructions. Briefly, a specific volume of extraction reagent from the kit was added to the muscle tissue, followed by homogenization and centrifugation to obtain the supernatant. A series of standard iron solutions were prepared. Blank, standard, and test tubes were set up: distilled water was added to blank tubes, standard solutions to standard tubes, and tissue supernatants to test tubes. Freshly prepared chromogenic reagent was then added to all tubes, which were incubated under specified conditions. After incubation, absorbance values at 593 nm were measured using a microplate reader. Iron concentrations in the muscle tissue samples were calculated based on absorbance values and the standard curve.

### Grip Strength Test

Maximal grip strength of mice was determined using a digital grip strength analyzer (Saiangsi Biological Technology Co., Ltd., China). Each mouse was placed on the force transducer platform and permitted to grasp the grid bars with all four paws. The experimenter then gently pulled the mice rearward by their tails at a consistent speed until grip release, with the mice maintained in a horizontal orientation throughout the entire procedure. Each mouse conducted five consecutive trials, with a 1-min rest period between each trial, and the highest value among the trials was used for subsequent data analysis.

### Rotarod Analysis

Before the formal test, mice were acclimated to the rotarod (Saiangsi Biological Technology Co. Ltd., China) for a 2-d period. The test initiated with a 5-min baseline phase at 5 revolutions per minute (rpm), followed by linear acceleration of rotational speed at 0.5 rpm/min until reaching 10 rpm; this speed was then maintained for 10 min. The final phase entailed continuous acceleration at 1 rpm/min until the mice became exhausted and fell off the rod. The total duration that each mouse remained on the rotarod was recorded as the primary outcome.

### Treadmill Endurance Test

Prior to the formal assessment, mice were acclimated to a treadmill (Saiangsi Biological Technology Co., Ltd., China) with a fixed 10° incline for 2 d. The treadmill endurance test comprised a single trial that began at 5 m/min for 5 min, followed by subsequent increments to 10 m/min for 10 min, 15 m/min for 15 min, and finally 20 m/min for 50 min. The test endpoint was defined as either accumulating 30 electrical shocks in total or remaining immobile on the shock grid for 2 consecutive seconds despite ongoing stimulation. The total distance traveled by each mouse during the test was documented.

### CatWalk Gait Analysis

Mice were acclimated to the CatWalk XT system (Noldus Information Technology BV, The Netherlands) for 7 d before the formal experiment. The walkway was calibrated with a 20 × 10 cm calibration sheet to convert pixel-based measurements to millimeters. Detection parameters were optimized through auto-detection, by adjusting the Camera Gain (10–20 dB) and Green Intensity Threshold (0.10–0.30) to eliminate background noise. During data collection, each mouse was placed at one end of the glass walkway (45 cm in length, designed for mice), and their runs were recorded at 100 frames per second until they entered the goal box. Runs were considered valid if they met the following criteria: a duration of 1–5 s, an average speed variation of ≤ 60%, and at least 2 valid runs per trial. Footprint classification was carried out using the Automatic Footprint Classification (AFC) function, with manual adjustments made for mislabeled footprints. Key gait parameters analyzed included mean intensity, print area, print length, print width, and maximum contact area.

### Histological Analysis

Muscle tissues were fixed in 4% paraformaldehyde, embedded in paraffin, and sectioned at 5 μm. H&E staining was performed to evaluate muscle morphology and fat infiltration. Nile red staining was used to detect lipid droplets in FAPs, with Hoechst 33258 staining for nuclei. Images were captured using a fluorescence microscope (Olympus BX53).

### Mouse Gastrocnemius Muscle Biopsy Processing for Flow Cytometry and Cell Sorting

Freshly isolated mouse gastrocnemius muscle biopsies were minced using fine dissecting scissors and digested at 37 °C for 60–90 min with gentle agitation. The digestion solution was prepared at a volume of 10 mL per gram of tissue, containing 2.5 U/mL Dispase II (Roche; catalog no. 4942078001), 1 mg/mL Collagenase II (Gibco; catalog no. 17101015), 5 mM MgCl₂ (Solarbio; catalog no. IM9034), and 2% penicillin-streptomycin (Gibco; catalog no. 15140122). Digestion was terminated by adding Phosphate-Buffered Saline (PBS) supplemented with 10% fetal bovine serum (FBS) and 2 mM Ethylenediaminetetraacetic acid (EDTA, Solarbio; catalog no. E1170). The resulting tissue suspension was sequentially filtered through 100-μm and 40-μm cell strainers (Falcon; catalog nos. 352360 and 352340, respectively) to obtain single-cell suspensions. After centrifugation and reconstitution, the cell concentration was adjusted to 2 × 10⁶ – 7.5 × 10⁶ cells/mL using FACS buffer (PBS with 2% FBS). Cells were then incubated with a fluorophore-conjugated antibody cocktail (details in Supplementary Table 1) for 30 min; this staining protocol targeted the identification of two key muscle-resident cell populations: SCs (phenotype: CD45⁻CD31⁻PDGFRα⁻ITGA7⁺) and FAPs (phenotype: CD45⁻CD31⁻PDGFRα⁺ITGA7⁻), with FAPs designated as the target population for sorting. Cell sorting and flow cytometric analysis were performed using a MA900 Multi-Application Cell Sorter, and all acquired data were analyzed with FlowJo software (version 10.4).

### LIP Detection

LIP was detected using the Calcein-AM fluorescent probe. Cells were incubated with 5 μM Calcein-AM for 30 min at 37 °C, and fluorescence intensity was measured using flow cytometry (SONY MA900). Higher fluorescence intensity indicates lower bioavailable iron.

### Intracellular Fe²⁺ Detection

Intracellular Fe²⁺ was detected using the FerroOrange fluorescent probe (Dojindo; Cat. No.: F374) in accordance with the manufacturer’s protocols. Cells were incubated with 1 μM FerroOrange working solution for 30 min at 37 °C with 5% CO₂. The medium was not replaced after incubation to avoid FerroOrange leakage. Fluorescence intensity was measured via flow cytometry (Model: SONY MA900; Channel: FL2). Higher fluorescence intensity indicates higher levels of intracellular free Fe²⁺.

### Transcriptomic Sequencing

Total RNA was extracted from purified FAPs, and its quality was assessed using an Agilent 2100 Bioanalyzer. The qualified RNA samples were subsequently subjected to library construction, which involved either mRNA enrichment via Oligo (dT)-coated beads or ribosomal RNA depletion, depending on experimental design. After RNA fragmentation, cDNA synthesis, and Illumina paired-end 150 (PE150) sequencing performed by Novogene (Beijing, China), raw sequencing data were filtered using Fastp to remove low-quality reads and adaptor sequences. The cleaned reads were aligned to the mouse reference genome (mm10) using the HISAT2 alignment tool, and gene expression levels were quantified with featureCounts. For differential expression analysis, DESeq2 or edgeR software was utilized to identify DEGs. The screening criteria for DEGs were set as |log₂FoldChange| ≥ 0.585 and P-value ≤ 0.05. Functional enrichment analysis of the identified DEGs was conducted using the clusterProfiler R package, with a focus on the “biological process” category within the Gene Ontology (GO) database. Enriched GO terms were considered statistically significant when the adjusted P-value (padj) was ≤ 0.05.

### RNA Isolation and qRT-PCR

Total RNA was isolated from FAPs via homogenization in TRIzol® Reagent (Invitrogen, Thermo Fisher Scientific, Waltham, MA, USA), with all procedures strictly following the manufacturer’s recommended protocol. For the determination of relative mRNA expression levels, the 2^-ΔΔCt method was employed, as reported previously.^[37]^ Cyclophilin A was selected as the internal reference gene to normalize the expression data and minimize sample-to-sample variation. All primer sequences utilized for qRT-PCR analysis are provided in Supplementary Table 2.

### Western Blot Analysis

Total protein was extracted from cells or muscle tissues using RIPA lysis buffer containing protease and phosphatase inhibitors. Protein concentration was determined using the BCA Protein Assay Kit. Proteins were separated by SDS-PAGE and transferred to PVDF membranes. Membranes were incubated with primary antibodies against HIF-1α, PPARγ, FTH1, NCOA4, and GAPDH overnight at 4 °C, followed by incubation with secondary antibodies for 1 h at room temperature. Bands were visualized using an ECL chemiluminescence system, and densitometric analysis was performed using ImageJ software. Details of the antibodies and related reagents are in Supplementary Table 3.

### Cell Culture and Treatment

FAPs were isolated from the skeletal muscles of mice via flow cytometric sorting and seeded into 96-well culture plates at a density of 5 × 10³ cells/well. The cells were cultured in DMEM/F12 medium (Acmec) supplemented with 10% (v/v) FBS and 1% (v/v) penicillin-streptomycin solution at 37 °C in a humidified atmosphere containing 5% CO₂, with daily medium replacement throughout the culture period.

For iron depletion treatment, FAPs were exposed to DFO (Solarbio) at final concentrations of 0, 1, 2, 5, 10, and 20 μM. For HIF-1α inhibition assays, FAPs were co-incubated with 2 μM PX478 (Solarbio) and 2 μM DFO. Cell viability was assessed using the CCK-8 assay kit (Solarbio) in strict accordance with the manufacturer’s protocols on day 3 and day 7 post-treatment, respectively.

Subsequently, FAPs were seeded into 12-well culture plates at a density of 2 × 10⁴ cells/well prior to adipogenic induction, which was performed using the OriCell® Mouse Bone Marrow Mesenchymal Stem Cell Adipogenic Differentiation Kit (Catalog No.: MUXMX-90031) following the manufacturer’s standardized instructions. After adipogenic induction, lipid droplet formation was evaluated via Nile Red-based fluorescent staining using the Lipid Fluorescent Staining Kit (Catalog No.: G1264; Solarbio) according to the manufacturer’s guidelines. Immediately after Nile Red staining, cells were incubated with 10 μg/mL Hoechst 33258 (Solarbio) in phosphate-buffered saline (PBS) at room temperature for 10–15 min in the dark to label cell nuclei. After staining, cells were washed 2–3 times with PBS to remove unbound dyes. All images were acquired using a Leica DMi8 fluorescence microscope, with simultaneous capture of Nile Red (lipid droplets) and Hoechst 33258 (cell nuclei) fluorescent signals.

### Statistical Analysis

Spearman correlation analysis and simple linear regression analyses were utilized to evaluate correlations among various parameters. Mann-Whitney U test and Kruskal-Wallis tests were employed for group-wise comparisons. Outliers exceeding the range of mean ± SD were excluded before data analysis. Statistical significance was defined as a two-tailed P value < 0.05, with P < 0.001 considered highly statistically significant. All data were expressed as mean ± SD. Statistical analyses were conducted with GraphPad Prism 9 software. Differences between two groups were analyzed using the unpaired t-test, while differences among multiple groups were assessed via one-way or two-way analysis of variance (ANOVA) followed by Tukey’s post-hoc test.

## Acknowledgments

This work was financially supported by the National Natural Science Foundation of China (Grant Nos. 82371957, 82502990); the Beijing Natural Science Foundation (Grant No. 7254373); the Beijing Physician Scientist Training Project (Grant No. BJPSTP-2024-08); the Beijing Jishuitan Research Funding (Grant No. JSTYC202207); the CMU Talent Project (Grant No. A2413); the Beijing Municipal Public Welfare Development and Reform Pilot Project for Medical Research Institutes (Grant Nos. JYY2023-8, JYY2023-11); the Beijing Municipal Health Commission (Grant No. BJRITO-RDP); and the Cultivation Project of Natural Science Foundation, Beijing Jishuitan Hospital, Capital Medical University (Grant No. ZR-202407).

## Author Contributions

Conceptualization: Q.R., R.W., L.W., and Y.L.

Methodology: Q.R., G.Y., D.W., and Y.W.

Investigation: Q.R., G.Y., W.W., J.F., K.M., A.G., M.F., Y.S., and Z.L.

Supervision: R.W., L.W., Y.L., and X.J.

Writing: Q.R., and R.W.

