## Supplemental Information for "Iron Deficiency Drives Sarcopenia in the Elderly: HIF-1α-Mediated Fibro-Adipogenic Progenitor Differentiation Induces Fat Infiltration and Impairs Muscle Function"

X. Jiang

Department of Trauma and Orthopedics, Beijing Jishuitan Hospital, Capital Medical University, National Center for Orthopaedics, Beijing, 100035, China.

L. Wang

Department of Radiology, Beijing Jishuitan Hospital, Capital Medical University, National Center for Orthopaedics, Beijing, 100035, China.

W. Wu, M. Fan, Y. Sun, Z. Lang, Y. Liu

Department of Spine Surgery, Beijing Jishuitan Hospital, Capital Medical University, National Center for Orthopaedics, Beijing, 100035, China.

*Corresponding authors

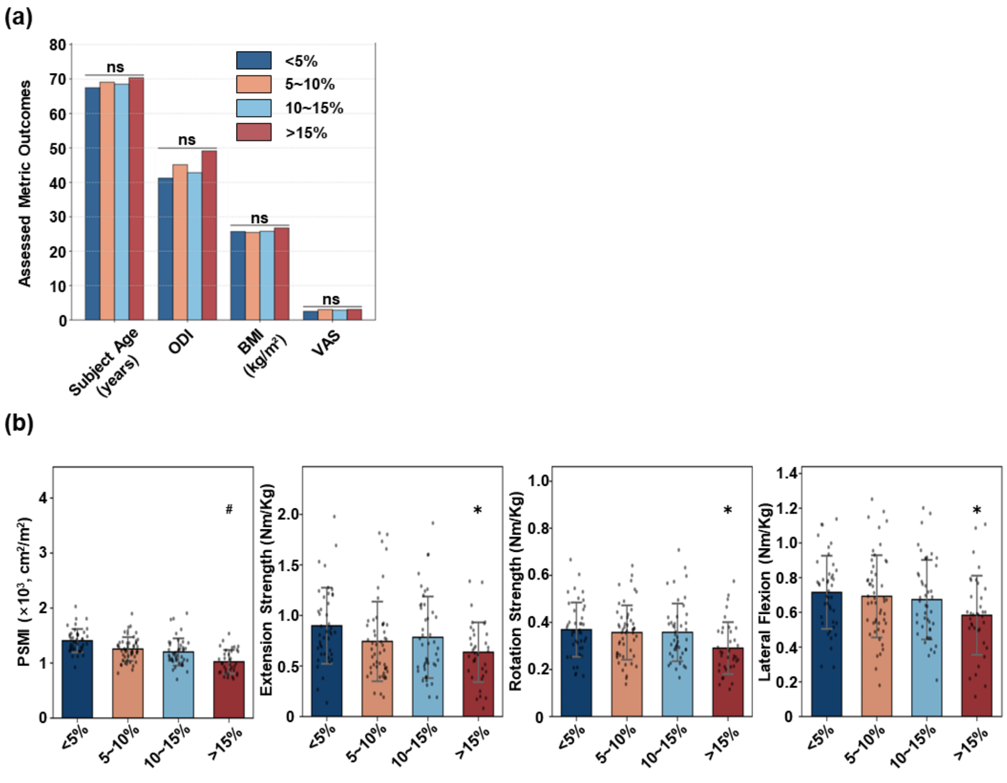


**Supplementary Fig. 1. Demographic, clinical, and muscle function characteristics across intramyocellular fat fraction (IMFF) subgroups.** (a) Demographic and clinical parameters (age, Oswestry Disability Index [ODI], body mass index [BMI], Visual Analog Scale [VAS] for low back pain) in participants stratified by IMFF subgroups (<5%, 5–10%, 10–15%, >15%). (b) Paraspinal muscle index (PSMI) and isokinetic muscle strength (extension, rotation, lateral flexion) across IMFF subgroups. Subgroup sample sizes: <5% (n=42), 5–10% (n=51), 10–15% (n=45), and >15% (n=36). Note: These baseline data represent the key findings of an independent study, which is currently under submission to *Advanced Science* (Manuscript ID: 8021953*)*. In the present study, we only use the IMFF subgroup stratification derived from these data to perform further analyses on other research objectives, and no overlapping core conclusions exist between the two studies. *Indicates a significant difference ($p < 0.05$) in the IMFF high subgroup compared with the low subgroup.

.

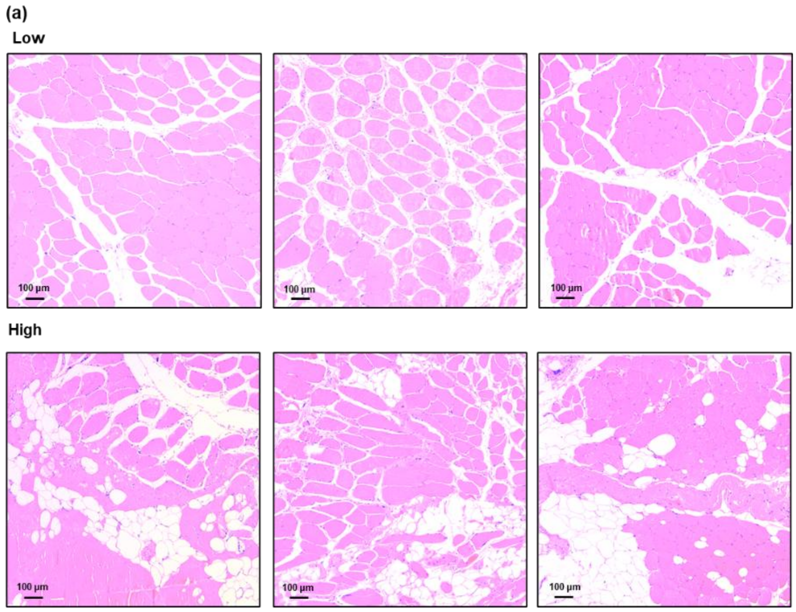


**Supplementary Fig. 2. Hematoxylin and eosin (H&E) staining of human paraspinal muscle samples illustrating distinct levels of fat infiltration.** (a) Representative H&E-stained sections of human paraspinal muscle tissues.

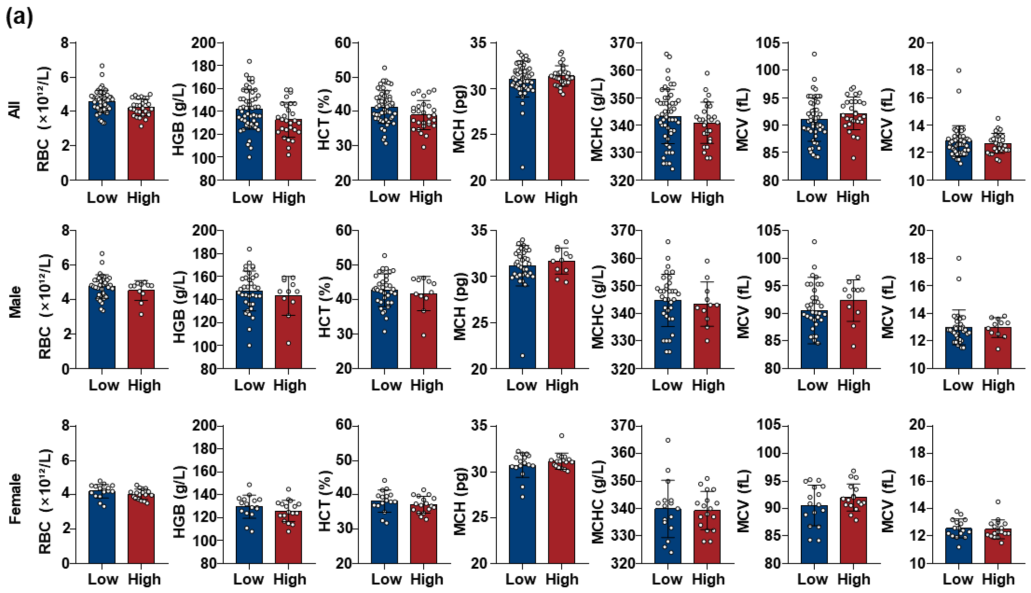


**Supplementary Fig. 3. Figure S3 Hematological parameters in Low vs. High IMFF groups.** (a) Comparison of anemia-related hematological parameters (hemoglobin [HGB], hematocrit [HCT], mean corpuscular volume [MCV], mean corpuscular hemoglobin [MCH], mean corpuscular hemoglobin concentration [MCHC], red blood cell count [RBC], white blood cell count [WBC]) between Low and High groups, stratified by gender (Low: n=51; High: n=28).

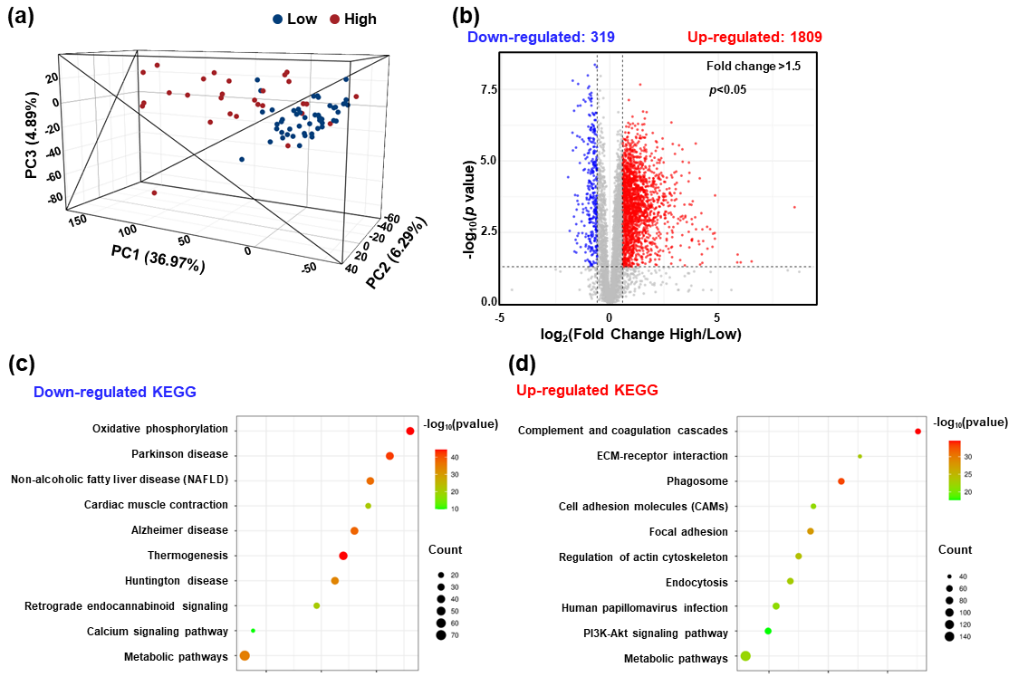


**Supplementary Fig. 4. Proteomic profiling and Kyoto Encyclopedia of Genes and Genomes (KEGG) pathway enrichment analysis.** (a) Principal Component Analysis (PCA) of proteomic data, showing distinct clustering of protein expression profiles between Low and High groups (PC1 = 36.97%, PC3 = 4.89%). (b) Volcano plot of differentially expressed proteins (DEPs) between High and Low groups. A total of 2128 DEPs were identified, including 1809 upregulated and 319 downregulated proteins ($|\log_2 \text{FoldChange}| > 0.585$, $p < 0.05$). (c) KEGG pathway enrichment analysis of downregulated DEPs. (d) KEGG pathway enrichment analysis of upregulated DEPs.

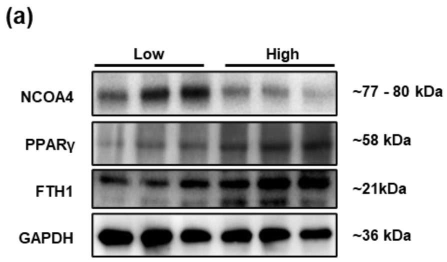


**Supplementary Fig. 5. Expression of ferritinophagy-related proteins in Low vs. High IMFF groups.** (a) Western blot analysis of NCOA4, PPARγ, and FTH1 in paraspinal muscle tissue from Low and High groups. GAPDH was used as an internal reference.

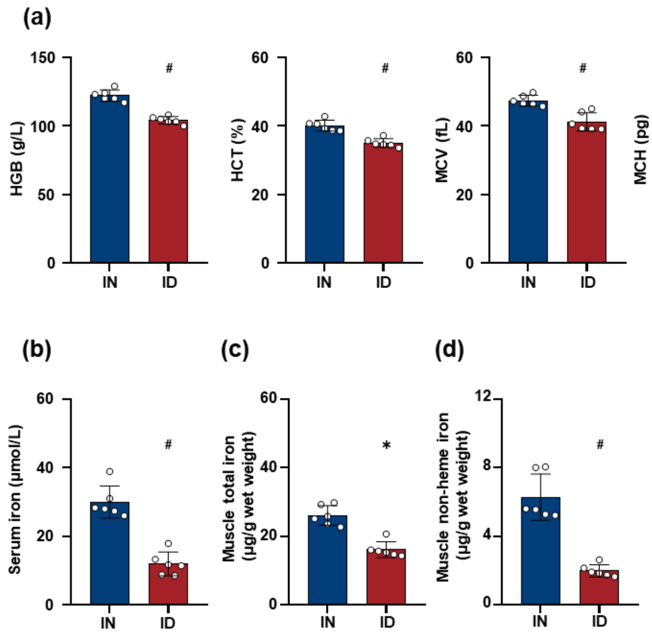


**Supplementary Fig.6. Validation of iron status in iron normal (IN) and iron deficiency (ID) models via hematological parameters and iron content analyses.** (a) Hematological parameters measured in IN and ID groups (n=6). (b) Serum iron concentration in IN and ID groups (n=5). (c) Muscle total iron content (μg/g wet weight) in IN and ID groups (n=6). (d) Muscle non-heme iron content (μg/g wet weight) in IN and ID groups (n=6). * and # indicate significant differences ($p < 0.05$ and $p < 0.001$) between IN and ID groups.

.

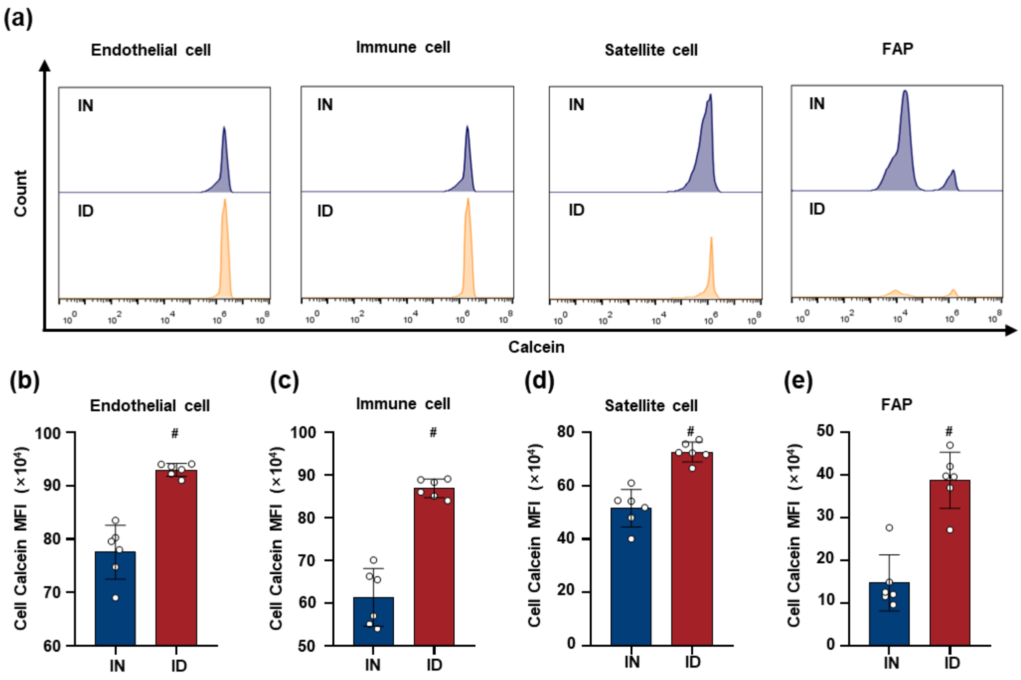


**Supplementary Fig.7. Flow cytometric analysis of Labile Iron Pool (LIP) in IN and ID mouse models.** (a–e) Analysis of LIP via Calcein fluorescence (higher fluorescence intensity indicates lower intracellular iron levels): (a) Histograms of Calcein fluorescence intensity for endothelial cells, immune cells, satellite cells (SCs), and fibro-adipogenic progenitors (FAPs) in IN (blue) and ID (orange) groups; (b–e) Calcein mean fluorescence intensity (MFI, $\times 10^4$) reflecting LIP in (b) endothelial cells, (c) inflammatory cells, (d)SCs, and (e) FAPs in IN and ID groups. * and # indicate significant differences ($p < 0.05$ and $p < 0.001$) between IN and ID groups.

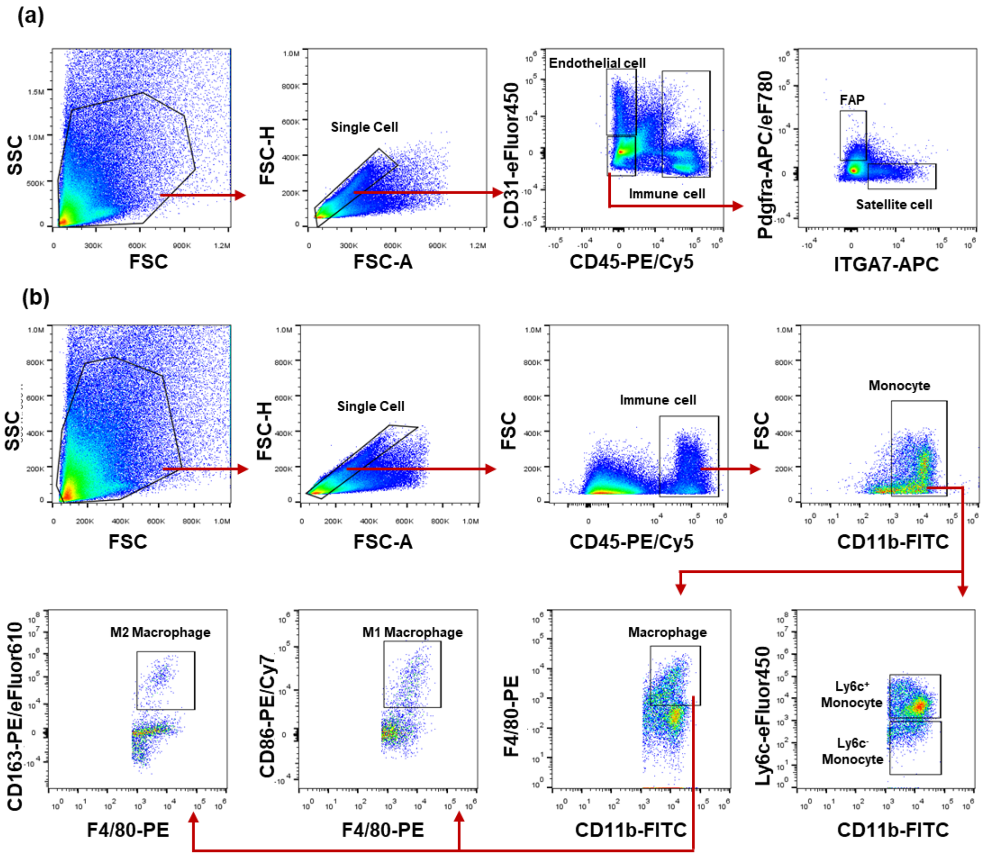


**Supplementary Fig.8. Gating strategies for flow cytometric analysis of cells in skeletal muscle single-cell suspensions.** (a) Gating for endothelial cells, inflammatory cells, SCs, and FAPs. (b) Gating for monocytes, Ly6c$^{+}$ monocytes, Ly6c$^{-}$ monocytes, macrophages, M1 macrophages, M2 macrophages and granulocytes.

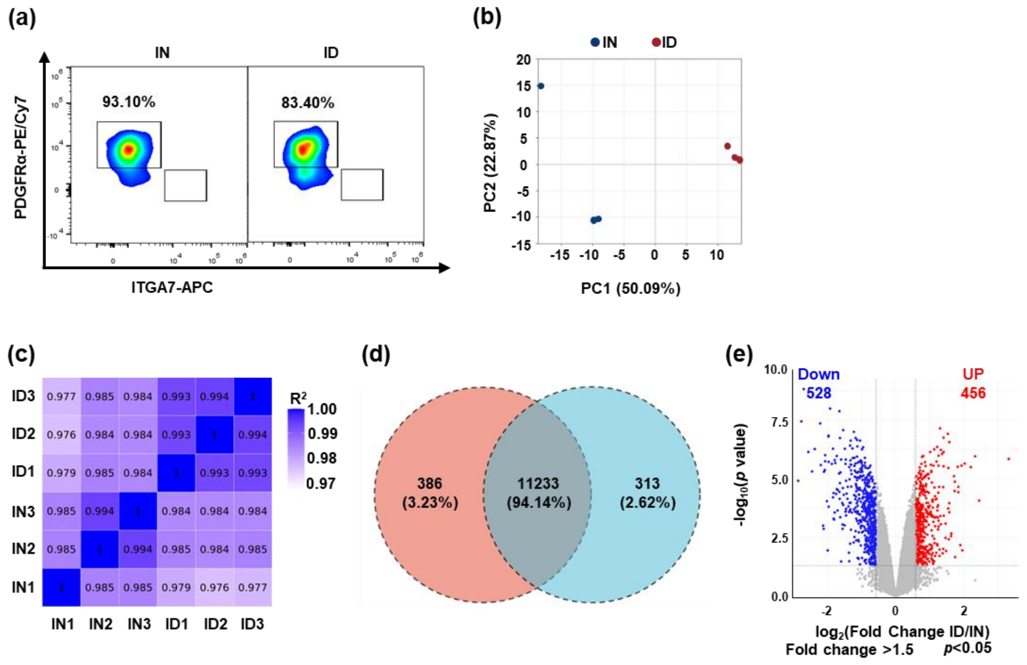


**Supplementary Fig.9. Validation of FAP purity and transcriptomic profiling of FAPs isolated from IN and ID mice.** (a) Flow cytometric verification of FAP purity: Dot plots (gated on PDGFRα-PE/Cy7 vs. ITGA7-APC) showing the purity of sorted FAPs (defined as PDGFRα$^+$ITGA7$^-$) in IN and ID groups; the respective purity percentages (93.10% for IN; 83.40% for ID) are labeled in the plots. (b) PCA of FAP transcriptomes: PCA plot (PC1 explains 50.09% of variance; PC2 explains 22.87% of variance) depicting the separation of FAP transcriptomic profiles between IN (blue dots) and ID (red dots) groups. (c) Correlation heatmap of FAP transcriptomic samples: Heatmap displaying $R^2$ (coefficient of determination) values among biological replicates of IN (IN1–IN3) and ID (ID1–ID3) groups; darker blue indicates higher transcriptomic similarity, demonstrating strong intra-group sample consistency. (d) Venn diagram of expressed genes in FAPs: Venn diagram illustrating the overlap and uniqueness of genes expressed in IN and ID FAPs; 11,233 genes (94.14%) are co-expressed, 386 genes (3.23%) are unique to IN FAPs, and 313 genes (2.62%) are unique to ID FAPs. (e) Volcano plot of DEGs in FAPs (ID vs. IN): Volcano plot identifying DEGs with screening criteria of $|\log_2(\text{Fold Change})| \geq 0.585$ (i.e., Fold Change > 1.5) and $p < 0.05$; blue dots represent downregulated genes (n=528) in ID FAPs relative to IN FAPs, and red dots represent upregulated genes (n=456).

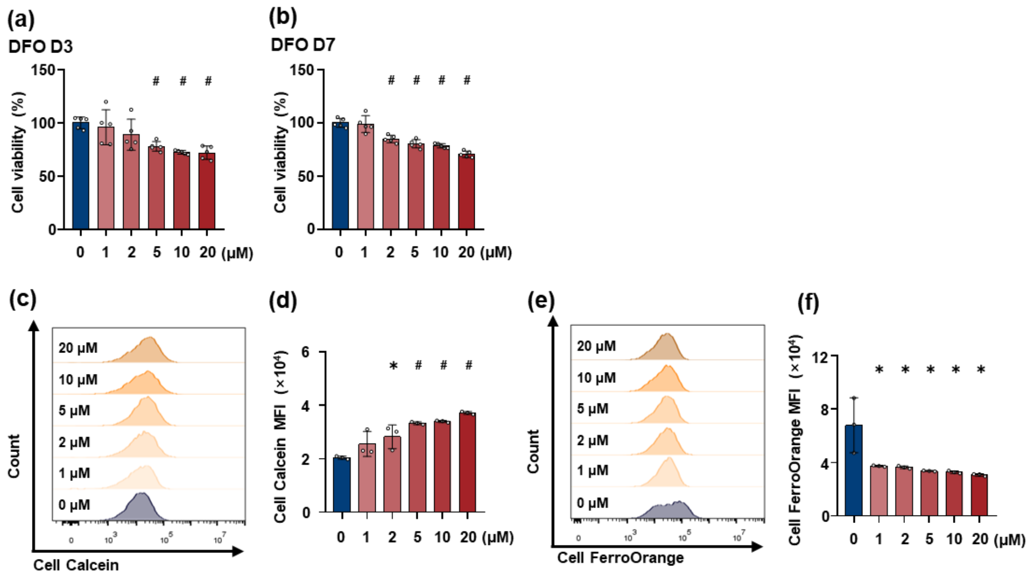


**Supplementary Fig.10. Cell viability and intracellular iron dynamics in FAPs treated with gradient concentrations of deferoxamine (DFO).** (a–b) Cell viability of DFO-exposed FAPs: Cell viability (quantified via CCK-8 assay) of FAPs treated with 0–20 μM DFO, measured at 3 days (D3, (a)) and 7 days (D7, (b)) post-treatment (n=5). (c–d) LIP detection via Calcein staining: Calcein fluorescence is quenched by intracellular iron, so reduce d fluorescence intensity corresponds to a higher LIP. (c) Representative fluorescence histograms of Calcein signals in FAPs treated with 0–20 μM DFO; (d) Quantitative analysis of Calcein mean fluorescence intensity (MFI, $\times 10^4$) corresponding to (c) (n=3). (e–f) Intracellular $Fe^{2+}$ detection via FerroOrange staining:FerroOrange fluorescence intensity directly reflects intracellular $Fe^{2+}$ levels. (e) Representative fluorescence histograms of FerroOrange signals in FAPs treated with 0–20 μM DFO; (f) Quantitative analysis of FerroOrange MFI ($\times 10^4$) corresponding to (e) (n=3). * indicates a significant difference ($p < 0.05$); # indicates a highly significant difference ($p < 0.001$) relative to the 0 μM (control) group.

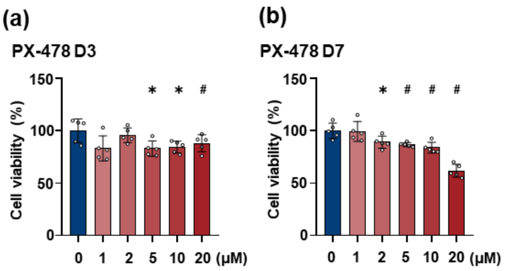


**Supplementary Fig.11. Effects of PX478 on FAP cell viability.** (a–b) Cell viability of FAPs treated with 0–20 μM PX478. Viability was measured at 3 days (D3, (a)) and 7 days (D7, (b)) post-treatment initiation. * indicates a significant difference ($p < 0.05$); # indicates a highly significant difference ($p < 0.001$) relative to the 0 μM group.

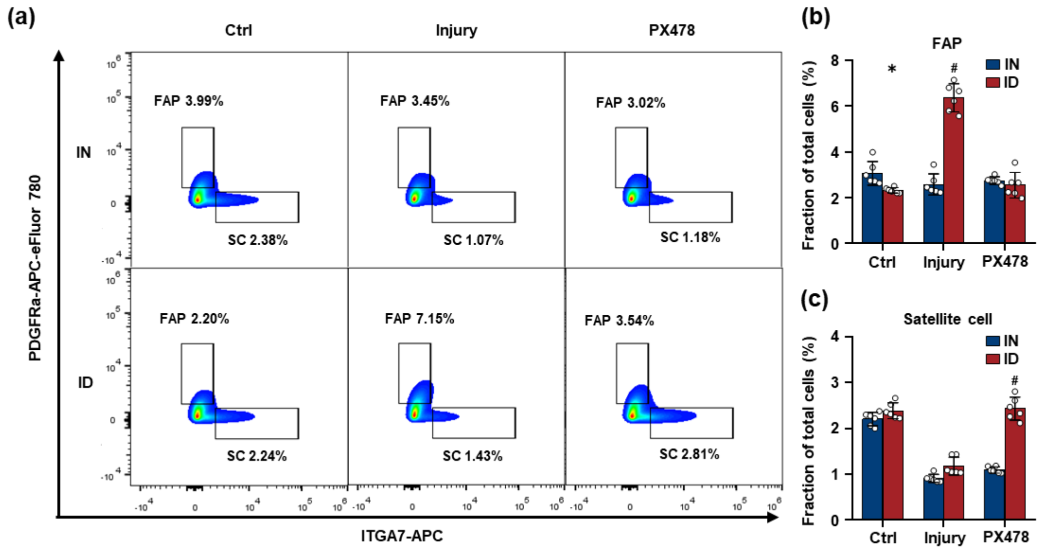


**Supplementary Fig.12. Flow cytometric analysis of FAPs and SCs in IN and ID models after glycerol-induced injury and HIF-1α inhibition by PX478.** (a) Dot plots depicting Dot plots depicting FAPs ($CD45^{-}CD31^{-}PDGFR\alpha^{+}ITGA7^{-}$) and SCs ($CD45^{-}CD31^{-}PDGFR\alpha^{-}ITGA7^{+}$) in Ctrl, Injury, and PX478 groups of IN and ID models, with respective population percentages labeled. (b) Fraction of FAPs among total cells in IN and ID groups across Ctrl, Injury, and PX478 conditions (n=6). (c) Fraction of SCs among total cells in IN and ID groups across Ctrl, Injury, and PX478 conditions (n=6). * and # indicate significant differences ($p < 0.05$, $p < 0.001$) between specified groups.

**Supplementary Table 1. Detailed information of fluorescent dye-conjugated antibodies used in flow cytometry.**

| Anti-mouse Ab | Fluorescent dye | Cat. No. | Manufacturer |
|---|---|---|---|
| CD45 | PerCP/Cyanine5.5 | 45-0451-82 | Thermo fisher |
| CD45 | PE-Cyanine5 | 15-0451-82 | Thermo fisher |
| CD31 | PE | 12-0311-82 | Thermo fisher |
| CD31 | eFluor™ 450 | 48-0311-82 | Thermo fisher |
| ITGA7 | APC | MA5-23555 | Thermo fisher |
| Pdgfrα | PE-Cyanine7 | 25-1401-82 | Thermo fisher |
| Pdgfrα | APC-eFluor™ 780 | 47-1401-82 | Thermo fisher |
| CD11b | FITC | 11-0112-82 | Thermo fisher |
| Ly6c | eFluor™ 450 | 48-5932-82 | Thermo fisher |
| F4/80 | PE | 12-4801-82 | Thermo fisher |
| CD163 | PE-eFluor™ 610 | 61-1631-82 | Thermo fisher |
| CD86 | PE-Cyanine7 | 25-0862-82 | Thermo fisher |

**Supplementary Table 2. Primers sequences used in this study.**

| Genes | Sequences (5'-3') |
|---|---|
| Pparg-F | TCGCTGATGCACTGCCTATG |
| Pparg-R | GAGAGGTCCACAGAGCTGATT |
| Adipoq-F | TGTTCCTCTTAATCCTGCCCA |
| Adipoq-R | CCAACCTGCACAAGTTCCCTT |
| Hmox1-F | AAGCCGAGAATGCTGAGTTCA |
| Hmox1-R | GCCGTGTAGATATGGTACAAGGA |
| Ccnb1-F | AAGGTGCCTGTGTGTGAACC |
| Ccnb1-R | GTCAGCCCCATCATCTGCG |
| Cdk1-F | AGAAGGTACTTACGGTGTGGT |
| Cdk1-R | GAGAGATTTCCCGAATTGCAGT |
| Bub1b-F | GAGGCGAGTGAAGCCATGT |
| Bub1b-R | TCCAGAGTAAAAGCGGATTTCAG |
| Cyclophilin A-F | AAGAAGGCATGAACATTGTGGAAGC |
| Cyclophilin A-R | CGGAAATGGTGATCTTCTTGCTGG |

**Supplementary Table 3. Western blot (WB) and Immunofluorescence (IF)-related antibodies and reagents used in this study.**

| Antibodies | Manufacturer | Cat. No. | Dilution |
|---|---|---|---|
| HIF1A | Thermo fisher | PA1-16601 | 1:1000 (WB) |
| HIF1A | Thermo fisher | PA1-16601 | 1:100 (IF) |
| PDGFRA | Thermo fisher | MA5-41209 | 1:1000 (WB) |
| PDGFRA | Thermo fisher | MA5-41209 | 1:100 (IF) |
| FTH1 | Thermo fisher | 701934 | 1:1000 |
| PPARG | Solarbio | K009215P | 1:1000 |
| NCOA4 | Proteintech | 10968-1-AP | 1:1000 |
| GAPDH | ZSGB-BIO | TA-08 | 1:1000 |
| Anti-rabbit IgG (H+L) | ZSGB-BIO | ZB-5301 | 1:10000 |
| Anti-mouse IgG (H+L) | ZSGB-BIO | ZB-5305 | 1:10000 |
| RIPA Lysis Buffer | Beyotime | P0013B | - |
| BCA Protein Assay Kit | Thermo fisher | A55864 | - |
| ECL Chemiluminescence Detection Kit | Epizyme | SQ201 | - |
| Lipid Fluorescent Staining Kit | Solarbio | G1264 | - |
| Hoechst 33258 | Yeasen | 40730ES03 | - |
